# Human-targeted drugs impact antibiotic sensitivity and growth of *Campylobacter jejuni*

**DOI:** 10.64898/2026.10.05.755907

**Authors:** Annamaria Zannoni, Roberto Olayo-Alarcon, Martin K. Amstalden, Medina Feldl, Rebecca Würz, Thea Bruchhardt, Johannes Kullmann, Sarah L. Svensson, Christoph Sotriffer, Mona Alzheimer, Christian L. Müller, Ana Rita Brochado, Cynthia M. Sharma

**Affiliations:** Department of Molecular Infection Biology II, Institute of Molecular Infection Biology (IMIB), Julius-Maximilians-Universität Würzburg; Department of Statistics, Ludwig-Maximilians-Universität München; Institute of Computational Biology, Helmholtz Munich; Department of Microbiology, Biocenter, Julius-Maximilians-Universität Würzburg; Institute of Pharmacy and Food Chemistry, Julius-Maximilians-Universität Würzburg; Interfaculty Institute of Microbiology and Infection Medicine Tübingen (IMIT), University of Tübingen; Center for Computational Mathematics, Flatiron Institute, New York; Cluster of Excellence ‘Controlling Microbes to Fight Infections’ (CMFI), University of Tübingen; Cluster for Nucleic Acid Sciences and Technologies – NUCLEATE, University of Würzburg

**Keywords:** *Campylobacter jejuni*, chemical genomics, screening, antibiotics sensitivity, non-antibiotics, efflux pump, antibacterials, stress response, promoter reporter, human targeted drugs

## Abstract

Antibiotic resistance is increasingly recognized as a major threat to global health, yet the contribution of non-antibiotics to this problem remains poorly defined. Here, we established a transcriptional chemical genomics approach to map how 2,415 host-relevant small molecules influence stress-responsive promoters and fitness of the foodborne pathogen *Campylobacter jejuni.* This screen uncovered widespread activation of the CmeABC efflux pump by diverse compounds, including several human-targeted drugs. Notably, ibuprofen, one of the most widely used non-steroidal anti-inflammatory drugs worldwide, induced efflux pump expression and thereby enhanced bacterial tolerance to ciprofloxacin. We also identified multiple commensal-sparing human-targeted drugs with growth-inhibitory activity against *C. jejuni*, highlighting opportunities for pathogen-targeted drug repurposing. Together, our findings reveal that non-antibiotic drugs can both promote resistance-associated phenotypes and selectively suppress pathogen growth. This underscores the need to consider collateral drug effects on bacterial physiology when designing strategies to curb the burden of antimicrobial resistance.

## Introduction

Antibiotic resistance is widely recognized as one of the most urgent global health challenges, spurring efforts to deepen our understanding of antimicrobial resistance (AMR) mechanisms and to develop new effective interventions. While antibiotics themselves have long been the focus, there is growing evidence that non-antibiotics and in particular human-targeted drugs can also perturb bacterial growth ^1,2^. While some drugs display broad antibacterial effects, others show some degree of species-specificity. Non-antibiotics can not only weaken host defenses against enteropathogens by altering microbiome composition and disrupting colonization resistance ^3^, but can also directly promote antibiotic resistance development in bacterial pathogens. For example, evidence in *Escherichia coli* indicates that non-antibiotics share similar resistance mechanisms to antibiotics ^1,4^.

Beyond antibiotics, bacterial pathogens are exposed to a variety of chemical stimuli during host colonization, which trigger stress response and virulence pathways critical for bacterial survival. For example, gut pathogens face different molecules derived from the host, diet, medication or the microbiome. Despite the potential risks posed by non-antibiotic drugs, we still lack a comprehensive landscape of which specific host chemicals are sensed by bacteria, what stress responses they elicit, and how these responses ultimately alter host-pathogen interactions, such as virulence and antibiotic resistance. To bridge this gap, transcriptional reporter libraries represent a powerful tool to study compound-mediated modulation of bacterial stress responses ^5,6^. Coupling specific stress responsive promoters to reporter genes enables simultaneous discovery of antibacterial compounds while providing insight into their underlying mechanisms of action and triggered pathways. For example in *E. coli,* both targeted approaches, such as using envelope stress reporters to identify compounds that perturb the Gram-negative cell envelope ^7^, or unbiased approaches that utilize genome-wide reporter libraries have been used to predict the mode of action of new compounds by comparing their transcriptional “fingerprints” to known antibiotics ^8,9^. Thus, we reasoned that screening a transcriptional reporter library in response to a library of host-related molecules is a powerful tool to identify chemical stimuli that trigger stress responses in bacterial pathogens. Moreover, considering variations in drug sensitivity across different bacterial species ^1^, different bacteria in the same niche likely respond to both shared but also species-specific chemical cues.

*Campylobacter jejuni* is the leading bacterial cause of foodborne gastroenteritis worldwide ^10^. Human infection is typically linked to consumption of contaminated poultry. Campylobacteriosis is characterized by diarrhoea, abdominal pain, and fever, but can also result in neurological sequelae such as the Guillain-Barré syndrome or even lead to life-threatening disease, particularly in young children, the elderly, or immunocompromised patients. While antibiotic treatment is reserved for severe cases, its efficacy is increasingly threatened by the rise of resistant strains. In particular, resistance to fluoroquinolones has reached critical levels in Europe, with ciprofloxacin being no longer recommended for the treatment of human infections^11^.

Despite its clinical relevance, the molecular mechanisms underlying virulence and stress resistance remain largely unknown in *Campylobacter*. Unlike other enteric bacteria, *C. jejuni* lacks many of the canonical global regulators and stress-defense sigma factors encoded by other pathogens (e.g., RpoS, RpoE) ^12,13^. Instead, it orchestrates stress adaptation and host-colonization via a restricted set of transcriptional regulators plus an expanding set of small RNAs (sRNAs), whose signals and triggers are poorly characterized ^14^. The reduced complexity of the *C. jejuni* regulatory architecture renders it a suitable model for dissecting stress responses elicited by non-antibiotic drugs, potentially facilitating the identification of key stress-response circuits that could remain hidden in organisms with higher redundancy. Moreover, it may increase *C. jejuni’s* vulnerability to chemical perturbations and illuminate unexplored properties of widely used pharmaceuticals, potentially informing drug repurposing strategies to mitigate the growing burden of AMR.

To profile *C. jejuni* stress responses, we assembled a transcriptional reporter library of genes involved in transcriptional and post-transcriptional control, stress responses, and antibiotic efflux, and screened it in a chemical-genomics high-throughput approach against 2,415 host-relevant compounds. In parallel, we explored the impact of these compounds on *C. jejuni* fitness. Our results reveal hundreds of chemicals that induce efflux pump expression and could concomitantly shift bacterial intrinsic sensitivity to antibiotics. Crucially, we demonstrate that the widely consumed and prescription free analgesic ibuprofen directly increases *C. jejuni* antibiotic resistance by disrupting efflux pump repression. Conversely, we identify commensal-sparing human-targeted drugs that inhibit *C. jejuni* growth, providing a candidate pool for pathogen directed drug repurposing strategies. We also developed a user-friendly interactive application to facilitate data exploration by the scientific community (link). In summary, our study provides an open-access resource for exploring chemical–genetic interactions in *Campylobacter* and provides new insights into how non-antibiotic compounds influence bacterial fitness and resistance-associated phenotypes.

## Results

### A transcriptional chemical-genomics screen uncovers *C. jejuni* stress response triggers

To decipher which environmental chemical stimuli are perceived by *C. jejuni* in the human gut and which stress responses they trigger, we designed a transcriptional chemical genomics screen. We exposed a promoter library of 26 transcriptional reporter strains to 2,415 chemically diverse host-related molecules at a single concentration (30 µM) in a high-throughput robotics platform and monitored bacterial growth and gene expression over time. In total, we profiled >60,000 compound–promoter pairs (**Fig. 1a**).

**Figure 1.**
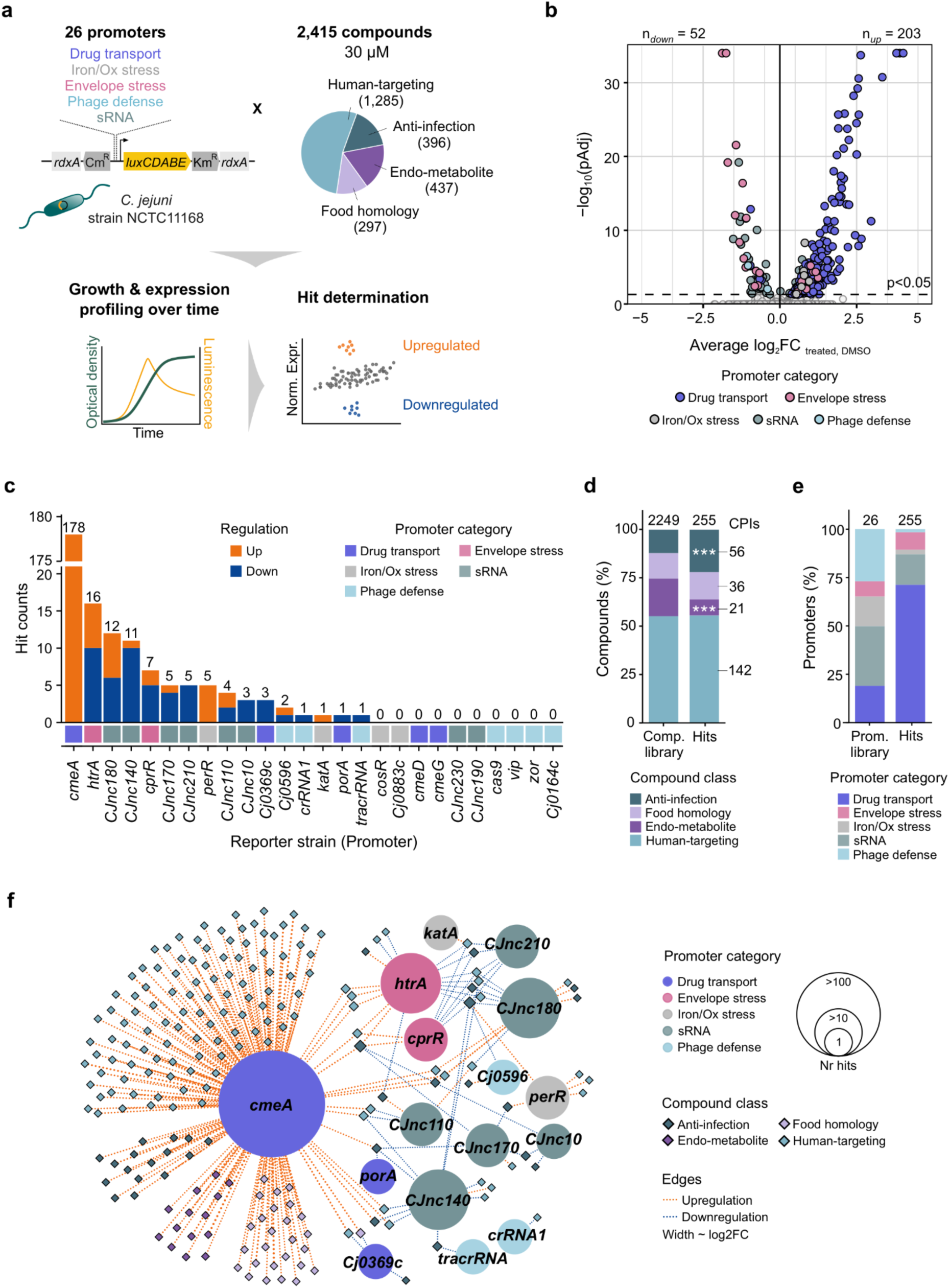
Unravelling stress responses triggered by host-relevant chemical stimuli in *C. jejuni*. **a)** Schematic overview of the transcriptional chemical genomics approach in *C. jejuni*. Lux-based transcriptional reporters of relevant stress response pathways (26 promoters) are challenged with a chemical library (2,415 compounds) mimicking chemical stressors faced by the bacterium in the host (human gut). Growth and luminescence were periodically measured over 22 h. Hits, i.e. compound-promoter interactions (CPIs), were cases of significant up- or downregulation of reporter expression upon compound treatment. **b)** Volcano plot of hits, displaying average log_2_ fold-changes (log_2_FC) across two biological replicates of normalized expression of promoters in the presence of compounds in comparison to average DMSO, plotted against -log_10_-transformed Benjamini Hochberg adjusted *p*-values with a cut-off of pAdj < 0.05. The highest pAdj value among the two replicates is used. Coloured circles represent hits according to promoter class, with the number of CPIs corresponding to up- and downregulation of promoter expression depicted above the volcano plot. For easier visualization, -log_10_(pAdj) was capped at 35. **c)** Hit count distribution for promoters separated into upregulated or downregulated. **d)** Proportion of compound classes in the screened library (2,249 excluding 166 strong growth inhibiting compounds) and in CPIs (255). Results of Fisher’s exact test for enrichment of compound classes among hit compounds compared to the screened library are indicated with asterisks. Significance: *** = pAdj < 0.001, * = pAdj <0.05. **e)** Proportion of promoter categories in the screened library (26 promoters) and in CPIs. **f)** Network representation of compound classes interacting with *C. jejuni* promoters. Promoters (circles) and compounds are displayed as nodes, colored by promoter categories and compound classes as in panel A. Edges represent the type of interaction (up- or down-regulations) colored as in panel C, and width represents log_2_FC.

The reporter library consisted of chromosomally integrated promoter–*luxCDABE* fusions in *C. jejuni* strain NCTC11168, enabling luminescence to serve as a proxy for promoter expression. We prioritized five major functional categories for promoter selection based on their relevance for *C. jejuni* survival in the gut and interaction with the host environment (**Fig. 1a, Suppl. Table S1**). This included promoters of (i) regulators and effectors of drug transport and AMR, such as a major porin and efflux pumps, which are critical for preventing intracellular accumulation of antibiotics and other toxic compounds (5 promoters); (ii) genes mediating oxidative and metal stress responses (4 promoters); (iii) envelope stress–related genes to monitor cell envelope interactions (2 promoters); (iv) sRNAs regulating different virulence traits of *C. jejuni* or with unknown function or stimuli (8 promoters); and (v) genes associated with anti-phage defense systems, including the Type II-C CRISPR-Cas system of *C. jejuni* ^15^ (7 promoters). A promoterless, but RBS-containing, *luxCDABE* strain was included as a control to measure background luminescence. The small-molecule library comprised 1681 Federal Drug Administration (FDA)-approved drugs, 297 food-derived compounds, and 437 endogenous metabolites (**Fig. 1a, Suppl. Table S2**). Of the FDA-approved drugs, nearly three-quarters (76.4%) were human-targeted drugs, while the remainder were anti-infective agents targeting bacteria, viruses, parasites, and fungi (**Suppl. Table S2**).

We periodically profiled growth (optical density at 600 nm) and promoter expression (luminescence) for all reporter strains over time in the presence of each compound (or DMSO as control) in biological duplicates. Growth was quantified as the baseline-corrected OD at 16 hours (OD_16h_; **Suppl Fig. S1a, left**), where most curves reached the onset of stationary phase, and promoter expression was integrated as the area under the luminescence curve until the same time-point (LuxAUC_16h_; **Suppl Fig. S1a, right**). *C. jejuni* growth in the absence of compound was consistent across all strains in this experimental setup (**Suppl Fig. S1b, left panel**), and highly reproducible between replicates (median Pearson, R = 0.84, **Suppl. Fig. S1c**). Promoter expression spanned a wide dynamic range (**Suppl. Fig. S1b, right panel**) with good replicate correlation for most strains (R = 0.80, **Suppl. Fig. S1c**). A subset of promoters (PCJnc230, P*vip*, P*zor*, P*cas9*) and the promoterless control (Δ*P-lux*) showed low Pearson correlation for LuxAUC_16h_ (R < 0.5), consistent with low signal-to-noise ratio, consequent to their inherently weak or absent promoter activity (**Suppl. Fig. S1c, S1d**). We excluded 5,170 non-concordant experiments, e.g., cases where technical errors occurred in one replicate and compounds reducing growth by more than 80% (166 compounds; **Suppl. Table S2, S3**) from the analysis. Overall, 60,599 compound-promoter pairs of the 26 promoter reporters and the empty promoter control were evaluated for significant expression changes (compound-promoter interactions, CPIs).

CPIs were identified by calculating the log_2_ fold-change (log_2_FC) of normalized expression (LuxAUC_16h_/OD_16h_) relative to the mean DMSO control for each compound-promoter pair, then applying a *z*-test against a promoter-specific DMSO null distribution to estimate *p*-values. A pair was called a CPI, or “hit”, if the highest FDR-corrected *p*-value across its two replicates was less than 0.05 (**Suppl. Table S4**). Overall, we identified 255 hits (CPIs, hit-rate 0.42%, **Fig. 1b, Suppl. Table S5**), with 203 and 52 up- and down-regulations, respectively. These 255 CPIs were driven by 207 unique compounds without significantly affecting growth (**Suppl. Fig. S2**), and while most compounds specifically triggered a single promoter, 28 triggered at least two (**Suppl. Fig. S3a**). Finally, CPIs were distributed across 16 out of 26 promoters (**Fig. 1c, Suppl. Table S6**). Promoters without CPIs may reflect low basal expression or absence of specific signals in our compound library for non-triggered promoters, e.g., phage-derived triggers (**Fig. 1c, Suppl. Table S6**).

### *C. jejuni* stress responses are largely triggered by human-targeted drugs

Compounds from all tested classes are represented within the CPIs (stressors, **Fig. 1d, Suppl. Fig. S3b**). Compared to their abundance in the chemical library, endo-metabolites were underrepresented among CPIs, while anti-infectives were enriched. The majority of anti-infective compounds among CPIs were, expectedly, conventional antibiotics, indicating that these molecules elicit bacterial stress responses even at sub-inhibitory concentrations. Beyond that, several anti-viral compounds were also triggers for *C. jejuni* promoters (**Suppl. Table S5**). Most strikingly though, *C. jejuni* stressors were predominantly human-targeted drugs (**Fig. 1d, Suppl. Fig. S3b**). With 142 out of 255 CPIs involving this compound class, drugs designed to interact with human targets, rather than microbial ones, accounted for nearly two-thirds of all stress triggers in our screen. This unexpected result establishes human-targeted drugs as a dominant source of chemically induced transcriptional stress response in our dataset.

More than 98% of the identified interactions were previously unknown for *C. jejuni* and ∼93% were, to our knowledge, unreported also for other bacterial species (**Suppl. Table S5**). Within these CPIs, the most responsive promoter class was drug transport, accounting for 71.4% of hits, followed by sRNAs with 15.9% of CPIs (**Fig. 1e**). The 255 hits of the 16 promoters showed varying numbers of interactions per promoter ranging from 1-178 (**Fig. 1c, 1f**). This showcases how promoters differed in sensitivity and specificity towards the profiled chemical cues. P*cmeA*, the promoter of the main *C. jejuni* efflux pump (CmeABC) was by far the most triggered promoter in our dataset, accounting for 67% of total interactions (**Fig. 1c, 1f**). For this, we were able to recapitulate known interactions, such as the induction of P*cmeA* by bile salts ^16^ or the efflux pump inhibitor Resveratrol ^17^. Beyond known *C. jejuni* CPIs, we also assessed some of our findings based on the biological features of the selected promoters and the literature. For example, we found some overlap between hits for the envelope stress reporters P*cprR* and P*htrA* (**Fig. 1f**), consistent with the CprR regulatory circuit, in which CprR positively regulates its own promoter and activates *htrA* ^18^. The second most responsive promoter class in our screen was sRNAs, with PCJnc180 and PCJnc140 accounting for more than half the hits in this class (**Fig. 1c**). This suggests they might participate in post-transcriptional regulation to adapt to diverse chemical stresses across distinct stress pathways. For example, our CPI network revealed partial hit overlap of CJnc180, which in concert with CJnc190 affects expression of the *C. jejuni* colonization factor PtmG ^19,20^, with the promoters of the efflux pump CmeABC as well as the envelope stress reporters HtrA and CprRS (**Fig. 1f**). Although further work is required to validate and investigate these compound-mediated regulations, our data can potentially expose previously unknown stimuli for *C. jejuni* sRNAs. Our dataset can be easily accessed and explored by the research community via our user-friendly interactive application (link).

These results establish our reporter library as a powerful tool for systematically mapping transcriptional chemical response in *C. jejuni*. Our screen reveals that *C. jejuni* stress responses are broadly responsive to chemical perturbation, in particular from non-antibiotics, with efflux and post-transcriptional regulation emerging as central adaptive mechanisms.

### NSAIDs induce promoter expression of the *cmeABC* efflux pump involved in AMR

Intrigued by the high number of hits for the P*cmeA* promoter and the critical role of the RND (resistance-nodulation-division) efflux pump CmeABC in conferring *C. jejuni*’s resistance to antibiotics, bile salts, and other antimicrobial agents ^16,21^ (**Fig. 2a**), we decided to focus on the chemical stimuli that impact *cmeABC* expression. Overall, 7.9% of all tested compounds in our screen upregulated P*cmeA* expression, corresponding to an approximately 18.8-fold enrichment over the overall screen hit rate, indicating that these compounds could affect *C. jejuni* susceptibility to antibiotics and other host-derived stresses. P*cmeA*-activating compounds spanned a wide range of chemical structures (**Fig. 2b**) and classes (**Fig. 2c**), suggesting that *C. jejuni* senses and responds to a broad spectrum of chemicals. Notably, only four pump activators were conventional antibiotics while the majority were human-targeted drugs with different therapeutic indications (**Fig. 2c**). Among these, NSAIDs were the most represented. We thus aimed to further investigate the connection between this class of compounds and its effect on P*cmeA*.

**Figure 2.**
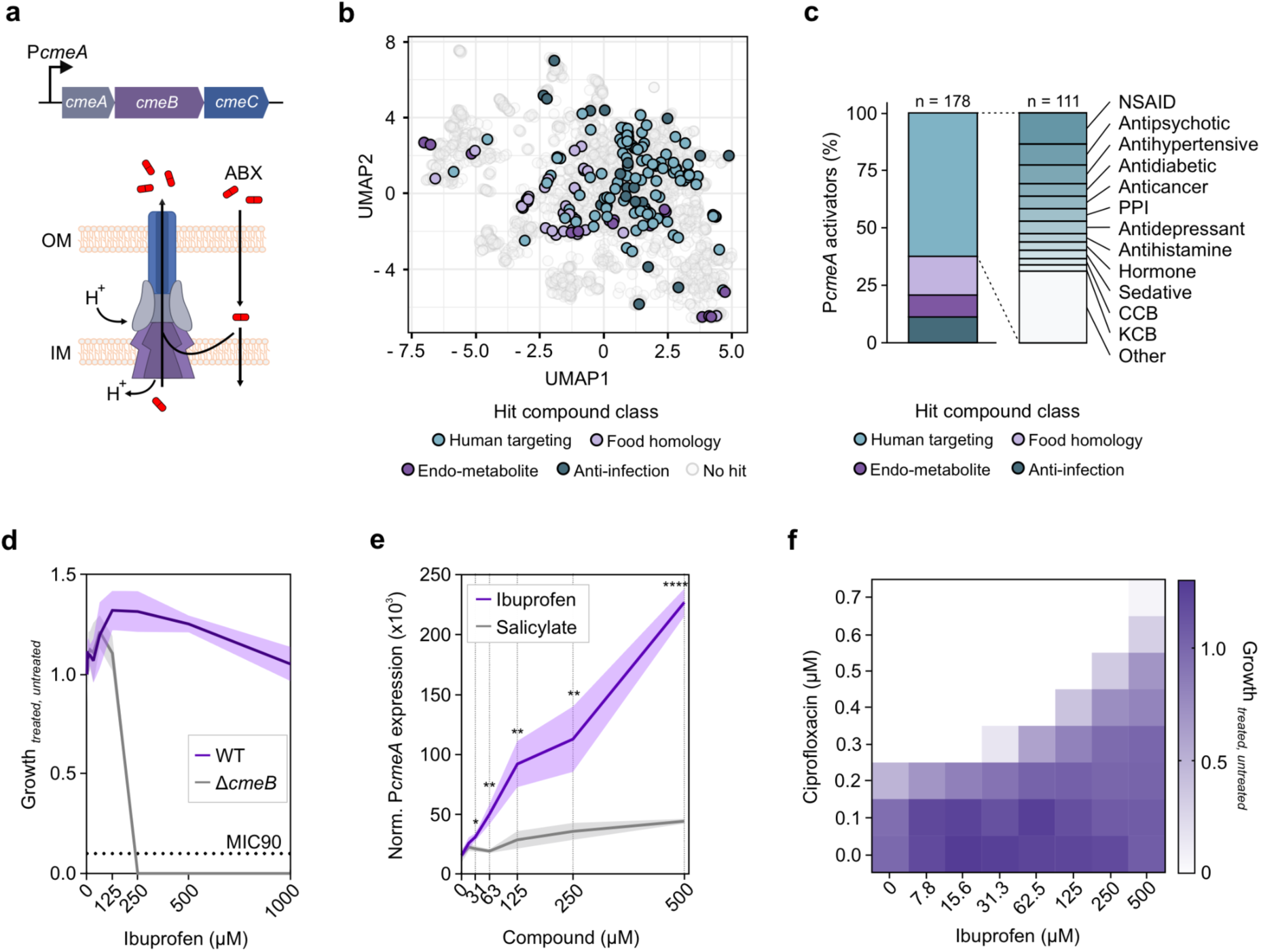
Human-targeted drugs stimulate efflux pump expression in *C. jejuni* and influence antibiotic resistance. **a)** Overview of the *cmeABC* operon, encoding for the RND-type CmeABC efflux pump, where *cmeB* encodes for the inner membrane (IM) component, *cmeA* for the periplasmic and *cmeC* for the outer membrane (OM) component. Cartoon depicts how CmeABC-mediated efflux confers *C. jejuni* resistance to toxic compounds, such as antibiotics (ABX) ^16,21^. **b)** Two-dimensional Uniform Manifold Approximation and Projection (UMAP) representation of chemical compounds tested in the screening, mapped according to Morgan fingerprints to reflect chemical structure similarity. P*cmeA* hits are highlighted and colored according to compound class. **c) Left:** Proportion of compounds activating P*cmeA*, divided by compound class as in panel B. **Right:** Proportion of therapeutic classes of human-targeting drugs that activate P*cmeA*. NSAID = non-steroidal anti-inflammatory drugs; PPI = proton pump inhibitors; EM = endo-metabolites; CCB = calcium-channel blockers; KCB = potassium channel blockers. **d)** Minimal inhibitory concentration (MIC) of ibuprofen tested in the concentration 0-1 mM, for *C. jejuni* WT and Δ*cmeB* strain. Data represent mean ± standard deviation (SD) for 3 biological replicates. Growth values in the presence of ibuprofen were obtained by dividing background corrected endpoint OD_600_ values upon drug treatment (treated) by the value of no-drug controls (untreated). **e)** Transcriptional reporter assay to compare dose-response activation of P*cmeA-lux* by the NSAIDs ibuprofen and salicylate, across a range of compound concentrations (31-500 µM). Data represent mean ± SD (n = 3 biological replicates) of growth normalized promoter expression in the absence (0 µM) or presence of the compounds. Results of Student’s *t*-test performed with GraphPad Prism comparing ibuprofen and salicylate treatments at the same concentration are indicated with asterisks. Significance: **** = *p*-value < 0.0001, ** = *p*-value < 0.01, * = *p*-value < 0.05. **f)** Checkerboard assay to assess the effect of ibuprofen and ciprofloxacin combinations on *C. jejuni* growth. Ibuprofen was tested in the concentration range of 0-500 µM and ciprofloxacin in the range of 0-0.7 µM. Growth values in the presence of single drugs and drug combinations were obtained by dividing background corrected endpoint OD_600_ values upon drug treatment (treated) by the value of no-drug controls (untreated). Data represent mean of three biological replicates.

NSAIDs showed a range of potency in activating P*cmeA* expression in our screen. Meclofenamate, meloxicam and tenoxicam were the strongest inducers, followed by (R)-(-)- ibuprofen (**Suppl. Fig. S4**). While the first three are widely used in treating arthritis, ibuprofen is an over-the-counter medication used to treat general pain and fever, typical symptoms of campylobacteriosis. As ibuprofen is one of the most commonly used drugs worldwide ^22^, we sought to further investigate this compound in relation to the *C. jejuni* efflux pump.

### Ibuprofen induces P*cmeA* expression and thereby antagonizes antibiotic sensitivity

Ibuprofen is commonly administered as a racemic mixture of R(-)-ibuprofen and S(+)-ibuprofen, and both enantiomers activated P*cmeA* in our screen (**Suppl. Fig. S4**), hence we decided to use a racemic mixture for our follow-up experiments. When examining the effect of ibuprofen on *C. jejuni* growth, we found that the bacterium is highly resistant to the drug, with a minimum inhibitory concentration (MIC) above 1 mM (**Fig. 2d**). Deleting the gene encoding the pump’s inner membrane component CmeB strongly reduced *C. jejuni* growth in the presence of ibuprofen, lowering the MIC to 250 µM (**Fig. 2d**). This suggests that the compound is a potential substrate of the efflux pump, as previously suggested for efflux pumps of other Gram-negative bacteria ^23^. To investigate the dose-response of the efflux pump promoter to ibuprofen, we tested the P*cmeA*-*lux* reporter strain against increasing compound concentrations. As a control, we included salicylate, a metabolite of the structurally related NSAID aspirin, which was previously shown to induce *cmeABC* expression and promote the emergence of fluoroquinolone-resistant *C. jejuni* ^24,25^. P*cmeA* indeed showed a dose-dependent response to both drugs in our transcriptional reporter assay (**Fig. 2e**). However, while the effect of our positive control salicylate was very mild, ibuprofen elicited a 5-fold stronger promoter response.

Since the CmeABC efflux pump reduces intracellular concentrations of antibiotics, we investigated whether the ibuprofen-induced transcriptional activation of the *cmeABC* promoter affected the sensitivity of *C. jejuni* to the fluoroquinolone ciprofloxacin, a known CmeABC substrate ^21^. To test this, we performed a checkerboard assay assessing *C. jejuni* growth in the presence of ciprofloxacin with increasing concentrations of ibuprofen. Remarkably, ibuprofen induced a concentration-dependent increase in the ciprofloxacin MIC of *C. jejuni* up to 2-fold increase in ciprofloxacin resistance (**Fig. 2f**). This effect was observed with physiologically relevant concentrations well within those reported for ibuprofen in the human small intestine ^26^. This finding reveals a strong antagonistic interaction between ibuprofen and ciprofloxacin, and suggests that co-exposure of *C. jejuni* to ibuprofen could strongly compromise antibiotic treatment efficacy.

### Ibuprofen induces efflux pump expression by alleviating CmeR repression

We next sought to uncover the molecular basis underlying the ibuprofen-induced expression of the *C. jejuni* efflux pump. CmeABC is under the control of CmeR and CosR, two repressors binding independently to two separate sites in P*cmeA* (**Fig. 3a**) ^27,28^. CosR is an essential orphan response regulator triggered by oxidative stress. CmeR is encoded immediately upstream of *cmeABC* and its flexible binding pockets have been shown to facilitate ligand binding, thereby inhibiting its repressor function ^24,25,29^. To investigate the possible involvement of the two regulators in the response to ibuprofen, we checked for differentially expressed target genes of CmeR/CosR upon ibuprofen treatment by RNA-seq (**Fig. 3b, Suppl. Table S7**). Together with the *cmeABC* operon, we identified 11 additional genes that were significantly upregulated in the presence of ibuprofen, and none of them are known targets of CosR ^27,30–33^. Likewise, the promoters of *katA* and *perR*, two CosR regulated genes, did not respond to ibuprofen or other P*cmeA*-activating compounds in our screen (**Fig. 1f**), further ruling out CosR involvement in the observed response to ibuprofen. Instead, we found two genes (*Cj0035c*, *Cj0561*) previously suggested to be under the control of CmeR ^34^ to be upregulated by ibuprofen. Our RNA-seq data thus clearly pointed to a CmeR-dependent derepression of *cmeABC* in the presence of ibuprofen.

**Figure 3.**
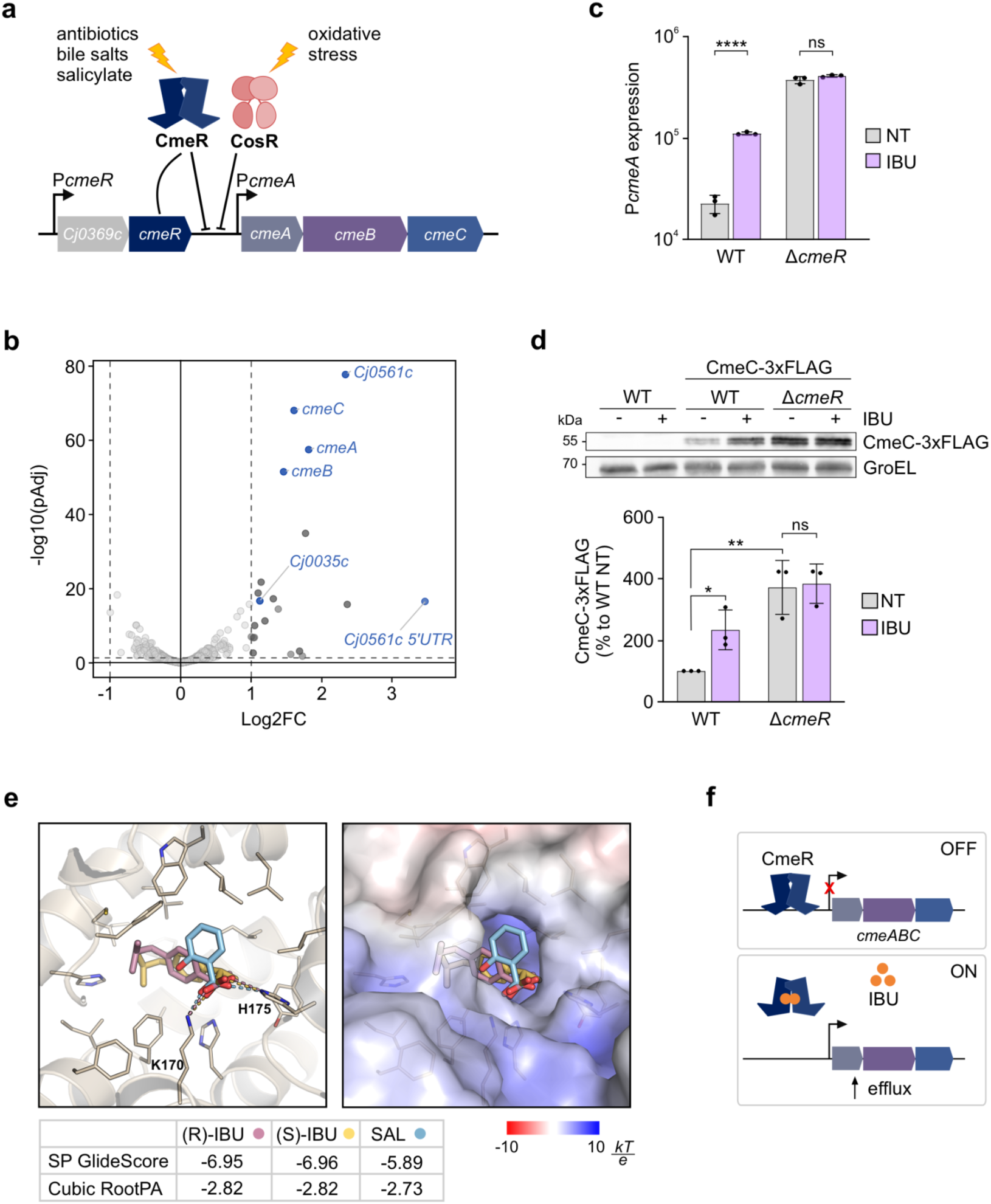
CmeABC activation by ibuprofen is mediated by CmeR. **a)** Schematic representation of known regulatory mechanisms of *cmeABC* expression. **b)** Volcano plot depicting differentially expressed genes after 30 min treatment with 150 µM ibuprofen identified by RNA-seq. Log2FoldChange (DESeq2, apeglm) in ibuprofen vs. DMSO treated samples (x-axis) is plotted versus –log₁₀ adjusted *p*-value (y-axis). The dashed horizontal line indicates the statistical significance threshold (FDR = 0.05), and vertical lines a log_2_FC of |1|. Significantly upregulated genes (log₂FC > 1, FDR < 0.05) are shown in dark grey or in blue if known CmeR targets, while non-significant genes are shown in light gray. **c)** Quantification of plate reader-based transcriptional reporter assay monitoring P*cmeA*-*lux* promoter responses in WT and Δ*cmeR* backgrounds, not treated (NT) or treated with 150 µM ibuprofen (IBU). The area under the curve (AUC) of luminescence curves was normalized by the AUC of growth curves (OD_600_). Data represent mean ± SD (n = 3 biological replicates). Results of Student’s *t*-test performed with GraphPad Prism comparing NT and IBU treatment are indicated with asterisks. Significance: **** = *p*-value < 0.0001, ns = not significant. **d)** Western blot analysis of protein samples of *C. jejuni* wildtype (WT) and 3xFLAG-tagged CmeC (CmeC-3xF) either in wildtype background or in Δ*cmeR* mutant strain. Samples were collected after 4 h of treatment with DMSO (NT) or 150 µM (IBU). CmeC-3XFLAG was detected with anti-FLAG antibody, and GroEL served as loading control. Protein levels were quantified with Fiji and are depicted as mean ± SD (n = 3 biological replicates). Results of Student’s *t*-test performed with GraphPad Prism comparing NT and IBU treatment are indicated with asterisks. Significance: ** = *p*-value < 0.01, * = *p*-value < 0.05, ns = not significant. **e)** Docking of NSAIDs IBU (both enantiomers) and salicylate (SAL) to CmeR, performed with the Glide module in Schrödinger Maestro. Left: top-scoring docking poses illustrating the suggested binding mode of the compounds to CmeR (PDB: 3QPS). Right: surface representation of the protein, colored by electrostatic potential (in units of kT/e). The ligands show good shape complementary in the binding pocket. SP GlideScores (more negative = more favorable) of the illustrated top-ranked docking poses are shown below, along with a size-normalized score to account for the different ligand volumes ^76^: Cubic Root PA = SP GlideScore/(number of heavy atoms)^1/3^. **f)** Model of proposed IBU-mediated regulation of *cmeABC* via CmeR. In basal conditions, CmeR binds to the *cmeABC* promoter region, thereby repressing its expression (OFF). Upon IBU binding, CmeR is released from the DNA and *cmeABC* expression is enabled (ON), leading to increased efflux of antibiotics.

To support the involvement of CmeR in this process, we used our transcriptional reporter fusion of P*cmeA* to compare promoter activity in WT and a Δ*cmeR* mutant upon treatment with ibuprofen. As expected, P*cmeA* activity in the WT was significantly increased upon ibuprofen treatment as well as upon deletion of the repressor (Δ*cmeR*) (**Fig. 3c**). However, no further upregulation was observed in the presence of ibuprofen in the Δ*cmeR* mutant. To assess the extent to which promoter activity correlated with protein levels, we 3xFLAG-tagged CmeC (encoded by the last gene of the *cmeABC* operon) to monitor protein level by western blot. Similar to promoter activity, CmeC protein levels were significantly induced by treatment with ibuprofen (**Fig. 3d**). While CmeC protein levels were generally upregulated by deletion of *cmeR*, treatment with ibuprofen did not increase protein levels further. These results indicate that ibuprofen activation of the efflux pump expression depends on CmeR (**Fig. 3c, 3d**).

To investigate whether ibuprofen could directly interact with CmeR, molecular docking simulations were performed using previously determined crystal structures of CmeR in complex with various bile salts ^29^. In addition to both ibuprofen enantiomers, we included the NSAID salicylate in the analysis, which was previously shown by electromobility shift assays to release CmeR from DNA ^24^. Docking predicted energetically favorable binding poses for both ibuprofen enantiomers and salicylate within the bile salt binding pocket of CmeR (**Fig. 3e**). In particular, (R/S)-ibuprofen and salicylate share the formation of hydrogen bonds to Lys170 and His175, while the lipophilic parts of the structures are placed in a hydrophobic region of the binding pocket. Similarly, the bile salts cholic acid and taurocholic acid engaged with CmeR Lys170 and His175 via hydrogen bonds in their crystallographic poses (**Suppl. Fig. S5**) ^24^. This suggests that ibuprofen could modulate CmeR activity through a similar mechanism as bile salts by binding to the same allosteric regulatory site.

Overall, our findings support a model where the exposure of *C. jejuni* to ibuprofen leads to a de-repression of its major efflux pump CmeABC, which is mediated by release of the CmeR repressor from the P*cmeA* promoter (**Fig. 3f**). Moreover, the upregulation of the efflux pump at the protein level in turn increases the intrinsic resistance of *C. jejuni* to the antibiotic ciprofloxacin.

### Identification of non-commensal targeting drugs that have bactericidal and bacteriostatic activity against *C. jejuni*

Besides transcriptional responses, our chemical genomics screen also monitored bacterial growth in response to compound treatment. Thus, we next focused on this readout to identify compounds that significantly impact *C. jejuni* growth and applied DGrowthR ^35^ to analyze differential growth responses in the presence of the compounds. We used the growth readout of the promoterless *C. jejuni* strain (Δ*P-lux*) upon treatment with the compound library (**Fig. 4a**). Here, we identified 10 compounds that increased and 207 compounds that decreased *C. jejuni* growth (**Fig. 4b, Suppl. Table S8**). While the growth promoting compounds were predominantly human-targeting drugs, growth inhibiting compounds were expectedly enriched for anti-infective agents with a particular strong representation of antibiotics from diverse classes (**Figure 4c, 4d upper panel**). However, we also found that antifungals (e.g., imidazoles) or artemisinin-derived antimalarials inhibited *C. jejuni* growth (**Suppl. Table S8**). Strikingly, however, 48.3% of growth-inhibiting compounds (n = 100) were not conventional antibiotics (**Fig. 4d, upper panel**), and nearly one-third of these (n = 29) had not previously been reported to affect bacterial growth (**Suppl. Fig. S6a**). The 100 compounds were predominantly human-targeted drugs (**Fig. 4d, middle panel**), including 21 compounds with potential antibacterial activity newly identified in our screen. To identify potential candidates with growth-inhibitory activity against *C. jejuni* while sparing gut commensals, we compared our findings with a previous study testing 1,197 compounds from the Prestwick Chemical Library against 40 representative gut bacterial strains ^1^. Of the 44 non-antibiotic *C. jejuni* inhibitors evaluated in both studies, 11 had minimal effect on the growth of gut commensals, highlighting their potential as selective agents for *C. jejuni*-targeted therapeutic approaches (**Suppl. Fig. 6b** and **c**, **Suppl. Table S8**). These included one anti-fungal, one natural product, and nine human targeted drugs from five different therapeutic classes. From the latter, we selected one representative compound per therapeutic class, choosing drugs that displayed varied growth-inhibitory potencies in our initial screen (**Fig. 4b, 4d lower panel**). These included the mucolytic bromhexine (BRO), a compound with reported growth inhibitory activity against *E. coli* and *Trueperella pyogenes* ^36^. We further selected the NSAID diflunisal (DIF) and the antihistamine clemastine (CLE), which were both reported to inhibit *Staphylococcus aureus* biofilms ^37,38^. Finally, we included the ADHD (attention-deficit/hyperactivity disorder) drug fipexide (FIP) and the antihypertensive Labetalol (LAB), both thus far without known effects against bacteria.

**Figure 4.**
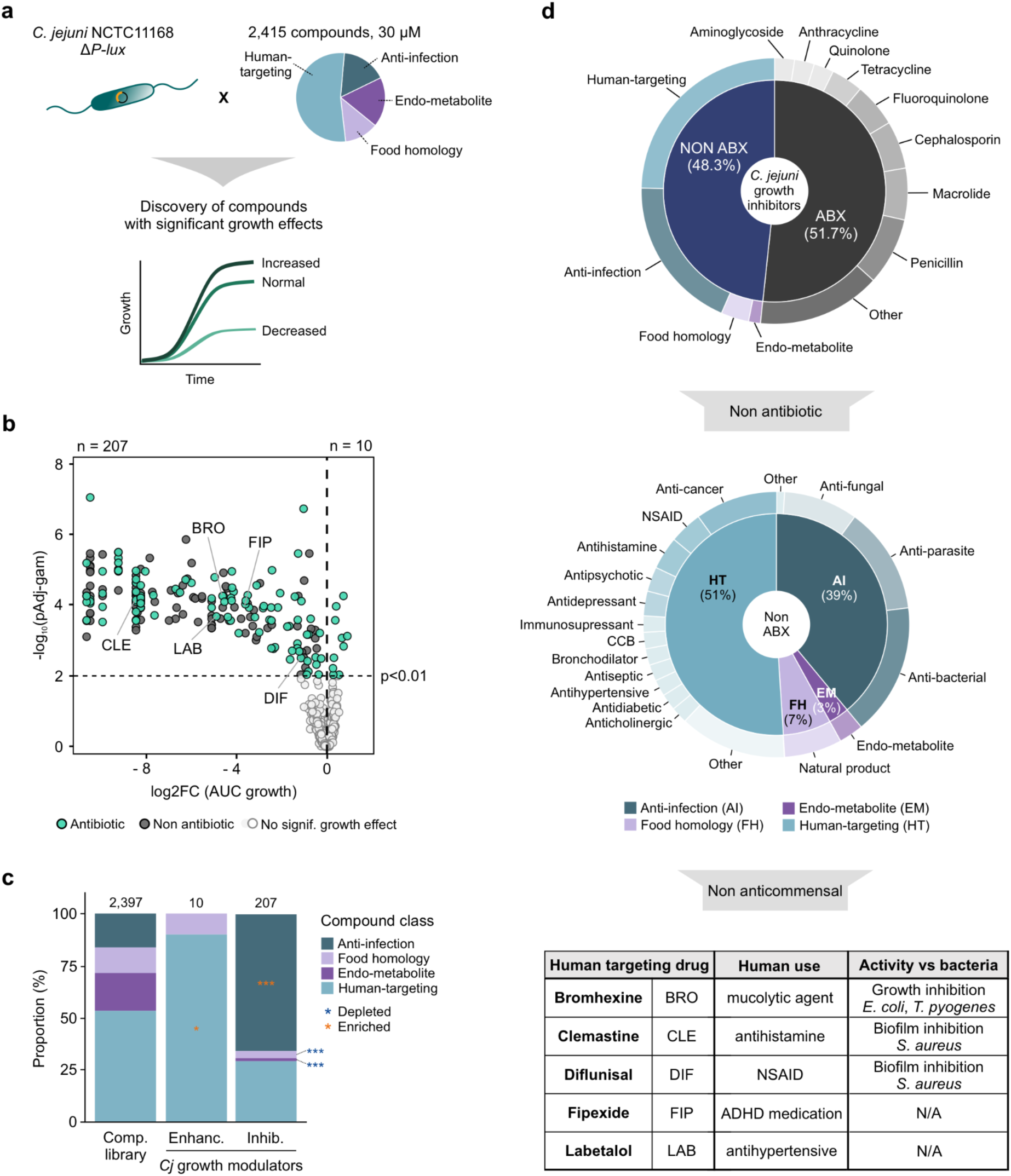
Discovery of compounds with significant effects on *C. jejuni* growth. **a)** Schematic depicting the identification of growth modulating compounds in *C. jejuni*. For this, growth readouts of the promoterless *C. jejuni* reporter strain (Δ*P-lux*) treated with 2,415 compounds were utilized. Legend: HT = human-targeted, AI = anti-infection, FH = food homology, EM = endo-metabolite. **b)** Volcano plot summarising the results of the statistically significant growth changes of Δ*P-lux* treated with 2,415 compounds. Corrected *p*-values (y-axis) are compared to overall growth changes (log_2_FC in AUC, x-axis). Growth modulating compounds are highlighted and colored based on their classification as antibiotic (green) or not (grey). **c)** Proportion of compound classes in the screened library (excluding three compounds with color that impacts growth assessment and 15 compounds excluded because of technical issues) and among *C. jejuni* growth enhancing (Enhanc.) and growth inhibitory (Inhib.) compounds. Compounds are colored according to classes. Results of Fisher’s exact test for enrichment of compound classes are indicated with asterisks. Significance: *** = pAdj < 0.001, * = pAdj < 0.05. Orange = enriched, blue = depleted. **d)** Selection process for compounds with diverse clinical applications, different growth-inhibitory potency against *C. jejuni* growth, and no anti-commensal activity. Upper: Donut plot of *C. jejuni* growth inhibiting compounds, divided between antibiotics (ABX) and non-antibiotics (NOT ABX). In the second layer, compounds are further split into sub-categories for non-antibiotics as well as antibiotics. Middle: Donut plot of non-antibiotic *C. jejuni* growth inhibitors, divided between compound classes (first layer) and therapeutic indication (second layer). Lower: five human targeted drugs (HTD) selected for further experiments, reported to have no growth inhibiting effect against gut commensal species, according to ^1^.

We first confirmed the growth-inhibitory activity of these compounds against *C. jejuni* in independent experiments and determined their MICs (**Fig. 5a**). To assess possible killing kinetics, we performed time-kill assays of exponentially growing *C. jejuni* exposed to 5x MIC of each compound (**Fig. 5b**). Whereas DIF, FIP, and LAB appeared to be bacteriostatic (maintaining constant CFUs/ml), BRO and CLE were bactericidal, with a 2-log10 and 4-log10 CFU reduction within 3 hours, respectively (**Fig. 5b**). Notably, both BRO and CLE reduced viable counts more than the antibiotic ciprofloxacin under the same conditions. To gain insight into physiological effects caused by the five selected compounds, we used confocal microscopy to reveal potential drug-induced morphological changes. Bacteriostatic DIF, FIP, and LAB caused no or only minor observable phenotypic changes compared to untreated cells (**Fig. 5c**). In contrast, bactericidal BRO and CLE induced pronounced DNA condensation into foci and reduced membrane staining. These two observations indicated that BRO and CLE could induce membrane damage, since membrane perturbation can trigger nucleoid compaction, a phenotype previously shown for some lipid biosynthesis inhibitors and membrane-active compounds in *E. coli* ^39^. For CLE, time-course imaging also revealed the formation of membrane protrusions in a subpopulation of cells over time (**Fig. 5d**), further prompting us to test whether BRO and CLE compromised outer membrane integrity. This was assessed in an uptake assay using the hydrophobic fluorescent dye *N*-phenyl-1-naphthylamine (NPN). Disruption of the outer membrane barrier facilitates NPN entry to the phospholipid layer of the inner membrane, resulting in a marked increase in fluorescence ^40^. We limited this analysis to CLE, due to non-specific interactions between BRO and the dye that prevented reliable assessment of membrane damage. We either left bacteria untreated as a negative control or treated them with ampicillin, which we hypothesized might not compromise outer membrane integrity in the short 10 min treatment window due to *C. jejuni*’s doubling time of ∼ 2 h. As expected, both controls did not increase fluorescence, indicating an uncompromised outer membrane (**Fig. 5e**). In contrast, treatment with the positive controls polymyxin B (a known outer-membrane permeabilizer) and EDTA (a chelator acting as membrane destabilizer) resulted in increased NPN fluorescence in a dose-dependent manner (**Fig. 5e**). Importantly, CLE induced an even stronger fluorescence increase than either of the positive controls. Altogether, these data indicate that CLE acts as a potent membrane-disrupting agent that leads to cell death of *C. jejuni*.

**Figure 5.**
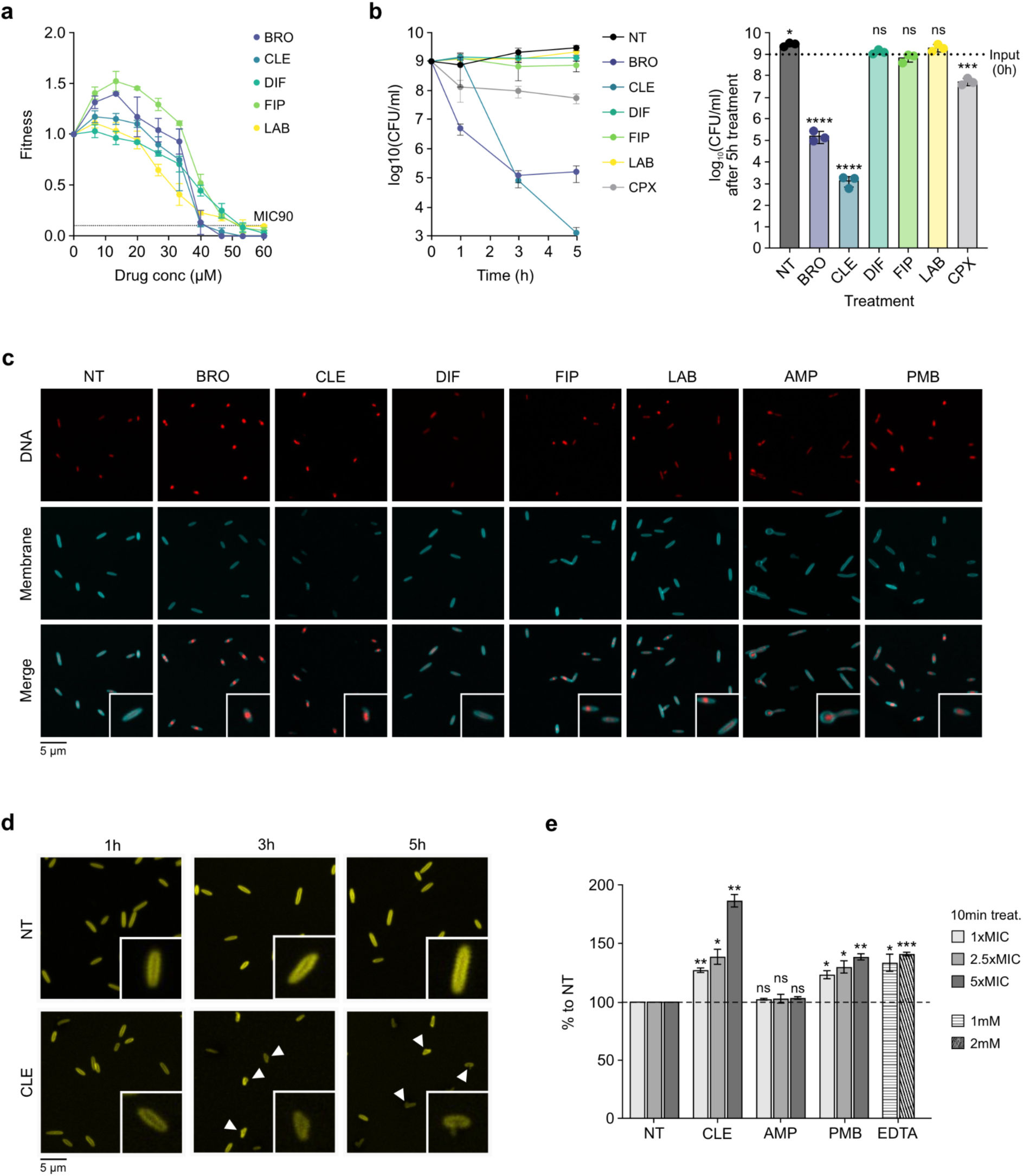
Drugs with non-bacterial targets show anti-bacterial activity. **a)** Fitness of *C. jejuni* WT grown in the presence of a concentration range (0-60 µM) of selected drugs. Threshold for Minimum Inhibitory Concentration required to inhibit 90% of bacterial growth (MIC90) is indicated. Data represent mean ± SD (n = 3 biological replicates). Fitness values in the presence of drugs were obtained by dividing OD_600_ values upon drug treatment by the value of no-drug controls at the same timepoint. Legend: BRO = bromhexine, CLE = clemastine, DIF = diflunisal, FIP = fipexide, LAB = labetalol. **b)** Time-kill assay of exponentially growing *C. jejuni* (10^9^ CFU/ml) treated over time with DMSO (NT) or with five times the MIC90 (5XMIC) of indicated drugs as in panel A or the antibiotic ciprofloxacin (CPX). Data represent mean ± SD (n = 3 biological replicates). Left: CFU quantification over time. Right: Comparison of viable CFUs after 5 h treatment with compounds, compared to untreated input (0h). Results of Student’s *t*-test performed with GraphPad Prism are indicated with asterisks. Significance: **** = *p*-value < 0.0001, *** = *p*-value < 0.001, * = *p*-value < 0.05, ns = not significant. **c)** Confocal microscopy of *C. jejuni* treated for 3 h with DMSO (NT) or 5XMIC of selected human-targeted drugs inhibiting *C. jejuni* growth as in panel A, or the antibiotics ampicillin (AMP) and polymyxin B (PMB). Bacterial membranes are stained with FM4-64 (cyan) and DNA with Hoechst (red). Images of NT, BRO, CLE, DIF, FIP, LAB are representative of three biological replicates and AMP and PMB treatments from one biological replicate, respectively. **d)** Confocal microscopy of *C. jejuni* treated over time with DMSO (NT) or with CLE at 5XMIC. Bacterial membranes are stained with FM4-64 (yellow). Images are representative of 3 biological replicates. Protrusions are marked by triangles. **e)** NPN-uptake assay to assess outer membrane damage, comparing cells untreated (NT) or treated for 10 min with CLE, negative control (-ctrl) antibiotic AMP, or positive controls (+ctrl) of membrane damage, e.g., the antibiotic PMB or the chelator EDTA at the displayed concentrations. MIC = MIC90. Data represent mean ± SD (n = 3 biological replicates). Significance vs. NT. *** = *p*-value < 0.001,** = *p*-value < 0.01, * = *p*-value < 0.05, ns = not significant (Welch’s *t*-test).

## Discussion

In this study, we generated the first resource mapping compound–promoter interactions in the widespread foodborne pathogen *C. jejuni*. By profiling 2,400 host-relevant small molecules, our dataset demonstrates an unprecedented variety of stressors that can trigger the expression of the bacterium’s CmeABC efflux pump. These include a number of NSAIDs, such as ibuprofen, for which we show how increased resistance to an antibiotic can arise as a collateral effect of efflux pump overexpression driven by a non-antibiotic compound. Furthermore, our screen revealed multiple commensal-sparing human-targeted drugs that inhibit growth of *C. jejuni*, highlighting opportunities for pathogen-targeted drug repurposing. By demonstrating that non-antibiotic compounds can affect gut bacterial growth as well as directly promote resistance-associated phenotypes in pathogens, our work highlights the need for a holistic understanding of antibiotic resistance beyond disruption of colonization resistance and genetic mutation. This broader mechanistic perspective may help inform the identification of new anti-infective strategies and deepen our understanding of AMR across diverse bacterial species.

By surveying an extended chemical library against 26 transcriptional reporters, our dataset complements the “PATHOgenex” resource ^41^, where *C. jejuni* whole transcriptome changes were profiled under 11 infection-relevant conditions. Our approach broadens this into a much larger chemical landscape and reveals the breadth of chemical stress sensed by *C. jejuni*. For example, 7.9% of our tested compounds upregulated P*cmeA*, suggesting that a substantial fraction of chemically diverse compounds could modulate *C. jejuni* susceptibility to antibiotics. Of these 178 compounds, only four were antibiotics, while the majority were human-targeted drugs, with NSAIDs emerging as the predominant class. A parallel screen in *S.* Typhimurium ^42^ found that inducers of the RND efflux pump AcrAB were also predominantly human-targeted drugs, although the specific compounds differed largely from those triggering CmeABC in *C. jejuni* (**Suppl. Fig. S7a**). Together with previous reports on efflux pump-inducing compounds ^43^, our data support a shared protective strategy across Gram-negative pathogens, in which RND-efflux pumps are upregulated in response to human-targeted drugs, while the inducing compounds appear to be species specific.

Moreover, approximately one third of the efflux pump activators identified in our study had previously been tested against a panel of gut commensals ^1^, and nearly half of them inhibited at least one out of 40 species (**Suppl. Table S5**). This suggests that CmeABC activation may help *C. jejuni* to better withstand compounds that inhibit gut commensals, offering a competitive advantage, as has been shown for other pathogenic bacteria, such as *E. coli* and *S*. Typhimurium ^3,44^. Conversely, 27.7% of *C. jejuni* non-antibiotic growth inhibitors identified in this study induce the *Salmonella* RND-efflux pump via the activator RamA, indicating specie-specificity in efflux pump responses and their potential role in protecting bacterial pathogens from chemical stress ^42^ (**Suppl. Fig. S7b**). Induction of RND-efflux pumps by non-antibiotics as a mechanism underlying protection from antibiotics has been also described for gut commensals, such as *Phocaeicola vulgatus*. For example, the anticoagulant drug dicoumarol selectively protects this commensal bacterium, but not pathogens, from antibiotics thus working as an “antidote” to minimize collateral damage by antibiotics in the gut ^45,46^. Overall, these findings indicate that presence and species-specific regulation of RND-efflux pumps by human-target drugs could be potentially relevant to inform targeted antibacterial therapies.

Since the protective agents broadly vary across bacteria, here we mainly focused on NSAIDs, and found that the commonly used over-the-counter drug ibuprofen induces CmeABC efflux pump expression, consequently increasing *C. jejuni r*esistance to the antibiotic ciprofloxacin. The aspirin metabolite salicylate has a similar effect in *C. jejuni* ^24^. Both compounds as well as additional NSAIDs also induce RND-efflux pumps in other Gram-negative bacteria, such as *E. coli* AcrAB ^23,47,48^ and *Pseudomonas aeruginosa* MexAB-OprM ^49,50^. However, efflux pump induction alone cannot predict the outcome of drug-drug interactions, as ibuprofen increases antibiotic resistance in both *C. jejuni* and *E. coli*, while it decreases it in *P. aeruginosa*. We further implicate CmeR, a TetR-family repressor, as the direct sensor of ibuprofen, analogous to the mechanism suggested for salicylate ^24^. On the other hand, ibuprofen-mediated stimulation of the *E. coli* AcrAB efflux pump depends on the MarA/MarR pathway, and salicylate is also an established MarR binder ^51^. This indicates a degree of functional overlap among the two different regulators. Overall, these findings open the possibility of designing species-selective therapies that exploit differences in bacterial regulatory networks to counteract the collateral effects of non-antibiotics that promote antibiotic resistance in bacteria.

In our screen, we also uncovered 91 novel non-antibiotic compounds, mainly human targeted drugs, that reduced *C. jejuni* fitness *in vitro*, substantially expanding (6.1x) the repertoire of compounds with growth inhibiting activity against this pathogen (**Fig 4d**, **Suppl. Fig. S6a, Suppl. Table S8**). Of these, 30 have, to our knowledge, not been described to affect bacterial growth in general (**Suppl. Fig. S6a, Suppl. Table S8**) or of *S.* Typhimurium in a parallel screen ^42^. Notably, 11 of the newly identified *C. jejuni* inhibiting compounds appeared to spare gut commensal species ^1^ suggesting they may preferentially target *C. jejuni* while preserving the surrounding microbial community. We validated five representative commensal-sparing human-targeted drugs with different therapeutic indications to be bacteriostatic (DIF, FIP, LAB) or bactericidal (BRO, CLE) against *C. jejuni*. While we could not find previous reports for FIP and LAB affecting bacterial virulence or growth, DIF was reported to reduce *S. aureus* biofilm formation by inhibiting the Agr system ^38,52^ and works as an “antidote” for the commensal *B. vulgatus* protecting it against the antibiotic doxycycline ^45^. Of the two bactericidal compounds, BRO was reported to directly inhibit *in vitro* growth of *E. coli* and *Trueperella pyogenes* and to synergize with diverse antibiotics ^36^. However, millimolar concentrations were required to achieve these effects. Notably, our microscopy experiments indicate that BRO might perturb *C. jejuni* membrane. Similarly to DIF, CLE also showed anti-virulence properties and inhibited *S. aureus* biofilms by targeting GdpP ^37^. However, it only had minimal effects on *S. aureus* growth at 200 µM, and similarly did not impact the growth of *S*. Typhimurium up to 150 µM ^42^. In addition, CLE was also reported to inhibit the growth of the intracellular pathogens *Coxiella burnetii*, *Legionella pneumophila*, and *Rickettsia conorii* in infected cells ^53^, and to potentiate macrophages against mycobacterial infection ^53,54^, although these effects were mostly believed to be host directed. Importantly, here we identify CLE as a potent antibacterial agent against *C. jejuni.* Our data suggest that its mechanism of action involves compromising the outer membrane integrity, thereby uniquely distinguishing it from all other mechanisms proposed for this drug. While the precise mechanism by which BRO and CLE might selectively target the *C. jejuni* membrane is unknown, we speculate that structural variations in the bacterial cell envelope may account for this specificity, analogous to how capsular polysaccharide differences in *E. coli* were recently shown to modulate sensitivity to certain antipsychotics ^55^.

Overall, our dataset provides a broad and open-access resource for understanding how *C. jejuni* senses and responds to host-relevant compounds. In particular, our findings show that non-antibiotic drugs can exert substantial and species-specific effects on bacterial physiology, including efflux activation and growth modulation, underscoring the need to consider the collateral effects of human-targeted drugs. At the same time, the identification of *C. jejuni* growth inhibitors across commensal-sparing compounds highlights the potential of this platform to guide novel pathogen-targeted therapeutic strategies.

## Materials and Methods

### Bacterial strains and culture conditions

Unless stated otherwise, *C. jejuni* strains (**Suppl. Table 9**) were routinely passaged on Müller-Hinton (MH) agar plates supplemented with 10 μg/ml vancomycin, or supplemented with the appropriate antibiotics for transformants selection: 20 μg/ml chloramphenicol (Cm), 50 μg/ml kanamycin (Km), or 250 μg/ml hygromycin B (Hyg). For liquid cultures, *C. jejuni* was grown in *Brucella* broth (BB) with shaking at 140 rpm at 37°C in a microaerobic atmosphere (10% CO_2_, 5% O_2_) without antibiotics.

### General recombinant DNA techniques

Plasmids and oligonucleotide primers (Sigma) used in this study are listed in **Suppl. Table 10 and 11**. For cloning, Phusion high-fidelity DNA polymerase was used (Thermo Fisher Scientific). Restriction enzymes, T4 DNA ligase and *Taq* polymerase for colony PCR (cPCR) were purchased from NEB.

### *C. jejuni* mutant construction

*C. jejuni* mutant strains (heterologous expression from the widely used *rdxA* locus, deletion, C-terminal 3×FLAG-tagging) were constructed by double-crossover homologous recombination with DNA fragments introduced by electroporation. Resistance cassettes used for cloning were either *aphA-3* (Km) ^56^, *aph(7’’)* (Hyg) ^57^, or *C. coli cat* (Cm) ^58^. Clones were validated for insertion of the desidered mutation by cPCR and verified by Sanger Sequencing (Microsynth, Macrogen).

Deletion mutants of genes encoding proteins were obtained by replacing their coding sequence with that of non-polar resistance cassette, by homologous recombination with a PCR-amplified DNA fragment. As an example, deletion of *cmeR* (Δ*cmeR*) was obtained by generating three PCR fragments: one, encompassing ∼500 bp of *Cj0368c* (*cmeR*) upstream region (CSO-2306/2307), a second one covering ∼500 bp downstream of *Cj0368c* (CSO-2308/2309) and a GmR cassette flanked by overlapping regions to the other two fragments (HPK1/HPK2). The overlap PCR product (CSO-2306/2309) was then used for *C. jejuni* transformation, and locus integration confirmed by colony PCR (cPCR) (CSO-2305/2857).

Epitope tagging at the C-terminal of coding genes was achieved by recombination with an PCR-generated DNA fragment inserting a sequence encoding 3xFLAG before the stop codon, followed by a non polar resistance cassette upstream of the native terminator sequence. For example, to produce the CmeC-3xFLAG strain, three PCR fragments were generated: a left flanking fragment encompassing ∼500 bp of *Cj0365c* until its penultimate codon (CSO-5071/5072), a right flanking region of similar length covering the region downstream of *Cj0365c* (CSO-5073/5074) and a partially overlapping middle fragment containing 3x-FLAG::KmR cassette (CSO-0065/HPK2). The fragments were fused by overlap PCR (CSO-5071/5074) and transformed into *C. jejuni* WT strain. FLAG tagging was confirmed by cPCR using CSO-5070/HPK2 and correct protein size was verified by western blot with Anti-FLAG antibody (see western blot section below).

### Cloning of transcriptional fusions for transcriptional chemical genomics screening

The *rdxA* locus (*Cj1066*) was used for heterologous gene expression in *C. jejuni* ^59^. To construct the promoter reporter library, a “promoter recipient strain” (CSS-7878) was first obtained by electroporation of *C. jejuni* WT with the plasmid pSSv166.6, constructed by subcloning into the previously constructed pSSv100.2 ^60(version^ ^1)^, which was modified to integrate the promoter of *Cj0367c* (P*cmeA*) upstream of the promoterless *hupB*(RBS)-*sfGFP* fusion, creating pSSh2.1. This was obtained by ligating a PCR fragment (P*cmeA*) amplified with CSO-5195/5196 on *C. jejuni* WT (CSS-5295) gDNA and the backbone of pSSv100.2, both digested with *Kpn*I/*Cla*I restriction enzymes. Finally, sfGFP was removed by amplifying pSSh2.1 with CSO-0762/5196, and swapped with the bacterial luciferase *luxCDABE* operon, amplified with CSO-5389/5477 on CSS-7048 (*E. coli* strain carrying pUT175-*luxCDABE* ^8^), after *Xma*I digestion of the PCR products and ligation. The obtained plasmid, named pSSv166.6, therefore contained the fragment *rdxA*::P*cmeA-hupB*(RBS)-*luxCDABE-KmR,* which was integrated in the *rdxA* locus of *C. jejuni* WT, obtaining CSS-7548. Genomic DNA was introduced into a clean *C. jejuni* WT background resulting in the “promoter recipient strain” strain CSS-7878. Each transcriptional reporter strain was then constructed by swapping the P*cmeA* promoter of CSS-7878 with an overlap PCR product containing the promoter region of the gene of interest, encompassing -200 to +10 nt from the annotated transcriptional start site ^14^, and a CmR cassette for selection in the promoter recipient strain at the *rdxA-lux* locus. As an example, the strain harboring the promoter of CJnc140 (CSS-7881), is described. First, we fused a 500 bp fragment of the *rdxA* downstream region (amplified with CSO-2277/5326 from *C. jejuni* WT gDNA) to a CmR cassette (amplified with CSO-5332/5333 from pGD34.7 ^60(version^ ^1)^) and an RBS-*luxC* fragment (amplified with CSO-5331/5330 from pSSv166) by overlap PCR with CSO-2277/5330. This fragment was ligated to pGEM-T (Promega) to generate plasmid pSSv165.3 (CSS-7380), which was validated by colony PCR with CSO-2277/5330 and Sanger sequencing with CSO-0644. The fused *rdxA*_DN-CmR fragment was then amplified with CSO-2277/5330 from this plasmid and introduced into *C. jejuni* CSS-7548 to generate strain CSS-7779, which was validated by colony PCR with CSO-0349/5330 and sequenced with CSO-0644. Next, a three-fragment overlap PCR was performed to fuse *rdxA*_DN-CmR with 1 kb of homology to *rdxA*, PCJnc140, and 1kb containing the RBS and part of *luxC*. The *rdxA*_DN-CmR region was amplified with CSO-5676/0574 from gDNA of CSS-7779, and the RBS- *luxC* region was amplified with CSO-5675/5637 from pSSv166. Finally, the CJnc140 promoter region was amplified with CSO-5665/5666 on *C. jejuni* WT gDNA to introduce regions of homology with the previous two fragments, and the three were fused by overlap PCR with CSO-5675/5676. The resulting fragment was used to electroporate CSS-7878 resulting in strain CSS-7880. For all strains generated using this approach, including CSS-7880, positive clones were validated by colony PCR with CSO-5682/5330 and Sanger sequencing with either CSO-5683 or CSO-5741. For the promoterless control strain used in the screening (Δ*P-lux*), the locus encompassing 1 kb of homology to *rdxA*, the CmR cassette and the promoterless but *hupB* ribosome binding site (RBS)-containing 1 kb *luxC* fragment was amplified with CSO-5675/5676 from CSS-7779 and transformed in the same background as the other reporter strains (CSS-7878), validated by colony PCR with CSO-5682/5330 and Sanger sequencing with CSO-5741.

### Transformation of *C. jejuni* by electroporation

Bacteria collected from passage one or two of *C. jejuni* strains grown on MH agar plates were resuspended in ice-cold electroporation buffer (272 mM sucrose, 15% (w/v) glycerol), pelleted by centrifugation (7000 x *g*, 5 min, 4°C), and washed twice. 50 μl of cell suspension were then mixed with 100-500 ng of purified DNA in a 1 mm-gap cuvette, electroporated at 2.5 kV for 4-6 ms in a Bio-Rad MicroPulser and immediately returned to ice. Cells were then mixed with 100 µL BB media and transferred to non-selective MH plates (supplemented with 10 μg/ml vancomycin), for overnight recovery at 37°C under microaerobic conditions. The following day, cells were transferred to selective MH agar plates containing the appropriate antibiotics and incubated until colonies appeared.

### High-throughput transcriptional chemical genomics screen

A custom chemical library of 2,415 small molecules dissolved in DMSO was assembled (MedChemExpress), including FDA-approved drugs, host metabolites, and food compounds. Multiple randomized wells containing only DMSO (18 wells/plate) served as untreated controls. Experiments were performed in 384 well plates (781903, Greiner Bio-One) and compounds (including control DMSO wells) were diluted to a final concentration of 30 μM in 40 µL BB using a Biomek i7 liquid handler (Beckman Coulter). *C. jejuni* transcriptional reporter strains precultures were grown overnight in 50 mL BB shaken at 140 rpm inside anaerobic jars (Cat. no. AG0025A, Oxoid) in the presence of CampyGen packs (Cat. no. CN025A, Thermo Scientific) for the generation of the microaerophilic environment. Screening plates were inoculated by pinning using a Singer Rotor+ replicator (Singer Instruments, UK). Plates were sealed with transparent breathable membranes (Breathe-Easy, Sigma-Aldrich-Merck) and incubated at 37°C, with 10% CO_2_, 5% O_2_ atmosphere, and shaking at 800 rpm using a Cytomat 2 incubator (Thermo Scientific). Growth (Optical Density at 600 nm, OD_600nm_) and transcriptional reporter activity (luminescence, Lux) were measured over 22 hours at 2 hours intervals in a Synergy H1 plate reader (Agilent, USA). The screen was performed in at least two biological replicates. Strains harboring promoters of putative anti-phage defense systems (P*vip*, P*zor*) and the respective upstream promoters (P*Cj0146c* and P*Cj0596*) were screened against a reduced number of compounds (1,725). In total, 65,703 promoter-compound combinations (pairs) were screened, including DMSO controls (3,258).

### Statistical workflow for hit determination

Data analysis was performed using R (version 4.2.2) and network visualization was done with Cytoscape 3.10.4. The data obtained from the high-throughput transcriptional chemical screening was analyzed using a custom workflow encompassing i) quality control, ii) expression value estimation, and iii) hit detection. These steps are briefly described below.

1. Quality control: Data quality was ensured by filtering experiments where technical failures occurred such as pipetting errors, abnormal growth or where sample contamination was observed (**Suppl. Table S3**). All retained growth curves were pre-processed with DgrowthR ^35^, removing the first 2 measurements (optical density at 600 nm, OD) and using the minimum of the 3^rd^ or 4^th^ OD measurement as baseline, subtracting its value from all subsequent measurements. Then, all OD measurements from a given library plate-replicate combination were scaled between the median OD value at the start and after 16 hours of growth (OD_16h_), corresponding to entry in the stationary phase for the majority of curves, measured for DMSO curves. We ensured that sufficient growth was achieved in each experiment by filtering experiments where the maximum OD measurement was lower than a data-driven threshold determined for each library plate-replicate combination. Experiments where compounds strongly affected growth were excluded, by applying a robust OD threshold specified as 20% of the 75^th^ percentile of OD_16h_ for DMSO curves (166 compounds, **Suppl. Table S2, S3**). Finally, we ensured that at least two replicates remained for each compound-promoter pair after all Åilters were applied. After applying these Åilters, 5,104 experiments were removed and 60,599 compound-promoter pairs remained (**Suppl. Table S3**).
2. Expression value estimation: the Åirst two timepoints of each luminescence (Lux) curve were ignored, consistent with growth data pre-processing. To quantify the Lux signal over time, we calculated the empirical area under the curve for each Lux curve (LuxAUC) up until 16 hours from the beginning of the experiment (LuxAUC_16h_). To account for the effect of the population size on the Lux signal, LuxAUC_16h_ was normalized by OD_16h_ as follows:

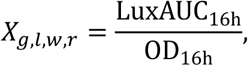

where *_X_*_g,*l*,w,*r*_ is the growth-normalized AUC Lux signal for a given promoter (*g*), plate (*l*), well (*w*), replicate (*r*) combination (promoter-compound-replicate experiment). In cases where more than two replicates for a given promoter-plate combination are present, we chose the two replicates that are most similar to each other by comparing the normalized Lux signal for all DMSO measurements of the promoter – library plate combination.

iii) Hit detection: To identify compounds that signiÅicantly up- or down-regulate promoter activity (compound promoter-interactions, CPIs, or hits), we Åirst estimate the fold change of each promoter-compound pair with respect to the average signal gathered from control experiments from exposure to DMSO:

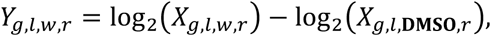

where *_Y_*_g,*l*,w,*r*_ is the log_2_ Fold Change (log_2_FC) of a given promoter-compound-replicate combination with respect to the average DMSO signal gathered from the same promoter-plate-replicate experiment (*_X_*_g,*l*,**DMSO**,*r*_). We determined that the log_2_FC values gathered from DMSO experiments followed a normal distribution in each promoter-plate-replicate combination by applying the Lilliefors test for normality. We next corrected these log_2_FC to account for non-specific compound and promoter effects. To do this, for each plate we construct a matrix:

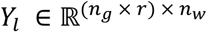

where *_n_*_g_ × *_r_* is the number of promoter-replicate combinations and *_n_*_w_ is the number of wells. We calculate a corrected fold change matrix by correcting each entry:

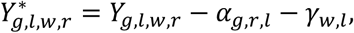

where *_α_*_g,*r*,*l*_ and *_γ_*_w,*l*_ are the promoter-replicate and compound effects, respectively. These effects are estimated by fitting an additive model using Tukey’s median polish procedure ^61^. In essence, row and column medians (*_α_*_g,*r*,*l*_ and *_γ_*_w,*l*_) are iteratively, removed until convergence criteria are met. In our work, we relied on the medpolish function from base R to accomplish this. Given that control experiments followed a normal distribution, we used these corrected fold change values to estimate z-scores for each promoter-compound-replicate combination ^(^*^z^*_g,*c*,*r*_^):^

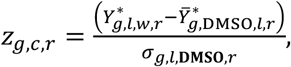

where *_Y_*^4 ∗^ and *σ*_g,*l*,**DMSO**,*r*_ are the average and standard deviation of the corrected fold change values gathered for DMSO experiments from the same plate as compound *c.* We then used these z-scores to estimate p-values following the z-test:

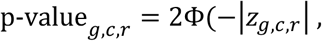

where Φ is the cumulative distribution function of the standard normal distribution. Finally, we retain the highest p-value for each promoter-compound pair and correct all retained *p*-values for multiple testing following the Benjamini-Hochberg method. We rely on a threshold of an adjusted *p*-value of 0.05 to identify hits (**Suppl. Table S4, S5**).

### Analysis of growth curves with DgrowthR

Growth curves were pre-processed and analyzed for differential growth phenotypes in response to chemical stressors using the R package DGrowthR ^35^. Briefly, each growth curve was baseline-corrected by setting the baseline to the minimum of the third and fourth OD measurements; the first two measurements were discarded due to technical noise. For the analysis of growth altering compounds performed on the promoterless strain (ΔP-*lux*), growth curves obtained in the presence of colored compounds (Methylene Blue, Evans Blue, and Tannic Acid) were excluded from differential growth analysis due to unreliable OD measurements. Additional 15 compounds were not considered for this analysis due to technical failure in one of the two replicates, as described in the quality control section of the pipeline of analysis for hits identification (**Suppl. Table S2, S3**). Differential growth analysis was performed by comparing growth curves collected in response to each chemical stressor to DMSO control curves from the same plate, using the DGrowthR’s multiple_comparisons function with 1,000 permutations. Significance was assessed using gamma-approximated p-values^62^, with a Benjamini–Hochberg adjusted p-value threshold of 0.01 (**Suppl. Table S8**).

### Low-dimensional representation of the chemical library

We gathered the SMILES for the compounds in the screening chemical library from PubChem using the pubchempy 1.0.4 Python module. SMILES were canonicalized and salts were removed using the rdkit 2020.09.1.0 python module. Binary Morgan fingerprints were calculated using rdkit, with parameters fp_radius=2 and fp_bits=1024. UMAP embeddings were computed with the R package uwot v0.2.4, using the jaccard distance matrix between compounds as input, with parameters n_neighbors = 20, n_components = 2, min_dist = 0.3. The jaccard distance matrix was computed using the native dist function in R, with method=”binary”.

### Total protein analysis by SDS-PAGE and western blotting

Bacterial cells grown to mid-exponential phase (OD_600_ of ∼0.4) were treated with compound at the indicated concentration for the indicated time, and harvested by centrifugation (13,000 rpm, 5 min, 4°C). The same volume of solvent (e.g. DMSO) was used for untreated conditions. Cell pellets were resuspended in 1× protein loading buffer (PLB; 62.5 mM Tris-HCl, pH 6.8, 100 mM DTT, 10% (v/v) glycerol, 2% (w/v) SDS, 0.01% (w/v) bromophenol blue) by shaking 5 min at 95°C. Samples corresponding to an OD_600_ of 0.1 were separated on 10% SDS-polyacrylamide (PAA) gels and blotted to a nitrocellulose membrane (semidry system, Peqlab). Membranes were blocked with with 5% (w/v) milk powder in TBS-T (Tris-buffered saline-Tween-20), and incubated overnight with primary antibodies at 4°C. Anti-FLAG (mouse monoclonal, 1:1,000; Sigma-Aldrich, #F1804-1MG) and anti-GroEL (loading control, rabbit polyclonal, 1:10,000, GE Healthcare, #RPN430) antibodies were prepared in 1X PBS. As secondary antibodies, either anti-mouse (sheep, 1:10,000; GE Healthcare, #RPN4201) or anti-rabbit (goat, 1:10,000; GE Healthcare, #RPN4301) polyclonal IgG conjugated with horseradish peroxidase were used, incubated in 3% BSA/TBS-T for 1 h at room temperature. Blots were developed using enhanced chemiluminescence reagent on a LAS-4000 Chemiluminescence Imager (GE Healthcare). A prestained Protein Marker (Thermo Fisher Scientific) was used for protein size estimation.

### Individual transcriptional reporter assays

Growth cultures were prepared in white, clear-bottom 96 well plates (Greiner BioOne) by diluting overnight cultures of *C. jejuni* reporter strains to a starting OD of 0.05 in 200 µL of BB. When appropriate, test compounds were also dissolved in the growth medium. For example, ibuprofen and salicylate were 2-fold serially diluted to obtain final concentrations between 0 (untreated) and 500 µM. Plates were sealed with breathable membranes and incubated at 37°C in microaerophilic atmosphere for 6 hours, with shaking at ∼750 rpm. Growth (optical density at 600 nm, OD) and luminescence (Lux) were measured with a plate reader (Infinite® M Plex, Tecan) in 1 hour intervals. To calculate growth normalized promoter expression, baseline correction was first applied to both raw OD and Lux curves for each individual sample well by subtracting the initial reading at 0 hours from each timepoint measurement. The area under the luminescence and growth curves (AUC) from 0 up to 6 hours was calculated using the trapezoidal rule (MESS R package), yielding AUC_Lux and AUC_OD respectively. For each of three biological replicates, normalized promoter activity was defined as the ratio between of cumulative luminescence to cumulative growth over the total time of the experiment (AUC_Lux/AUC_OD).

### Docking of small molecules to CmeR

The available crystal structures of CmeR (3QQA (2.20 Å resolution), 3QPS (2.35 Å), 3HGG (2.57 Å), 3HGY (2.42 Å), 2QCO (2.25 Å)) were retrieved as biological assemblies from the PDB (Protein Data Bank at www.rcsb.org). Structure preparations were carried out with Maestro 2023-4 (Schrödinger, LLC) using the *protein preparation workflow*. Hydrogens were added corresponding to the protonation state at pH 8.0, while terminal residues of the polypeptide chains were capped. Ligands were protonated for a pH of 8.0 using *LigPrep* with *Epik* ^63,64^. The OPLS4 force field was used for geometry optimization using default settings ^65^. Docking was performed with *Glide* ^66,67^ through maestro 2023-4 using the *SP* and *XP* docking modes. Before grid generation, solvent molecules of the crystal structure were removed. The binding sites were defined by the centroid of the native ligand with an inner box size of 10 Å (cubic edge length) and an outer box size of 23 Å. The docking protocol was optimized by a re-docking study. Based on the results, PDB 3QPS was selected as the target structure for the prospective docking. Ligand sampling was carried out by including input ring conformations, penalizing non-planar amide conformations, and enhancing planarity of conjugated pi-systems. All further settings were kept at their default values. 500 poses were included for post-docking minimization, and the number of output poses was limited to 50. Docking poses of a given ligand were hierarchically clustered with cut-off of 1.00 Å and sorted by the *Emodel* score. Figures of molecular structures were generated with Pymol 2.6.1 (Schrödinger, LLC). The electrostatic potential mapped to the surface was calculated with the APBS Electrostatics Plugin ^68^ using default settings (0.15 M ionic strength in monovalent salt, 310 K, protein dielectric 2, and solvent dielectric 78). Radii and charge assignment was performed with PDB2PQR ^69^ at pH 8.0 using Amber parameters.

### MIC determination for compounds with growth-inhibitory activity

Non-antibiotic compounds were diluted linearly to obtain 9 equally spaced concentrations starting from 60 µM, while antibiotics were 2-fold serially diluted 9 times, starting from 40 µM to 0 µM. Experiments were conducted in 96-well plates (Greiner BioOne), with a total volume of 200 µl, and inoculated with a starting OD_600_ = 0.001 of *C. jejuni* (∼1 x 10^6^ CFU/ml, corresponding to ∼2 x 10^5^ bacteria per well) from an overnight culture. Plates were sealed with breathable membranes and incubated at 37°C in microaerophilic atmosphere for 23 hours, with shaking at ∼750 rpm. OD_600_ was measured with a plate reader (Infinite® M Plex, Tecan) and end-point growth across drug concentrations was calculated for each well by subtracting OD_600_ values at 23 hours and at 1 hour, to account for both well-specific background as well as membrane clarity. The Minimal Inhibitory Concentration (MIC) was considered as the concentration at which growth was inhibited more than 90% compared to non-treated controls (MIC90). As calculations for drug inhibitory concentration are known to have a twofold error margin ^70^, we considered MIC within 15-60 μM as being in agreement with the screening results.

### Checkerboard Assay

Checkerboard assay was performed to assess the effect of the combination of the antibiotic Ciprofloxacin (CPX) and the non-antibiotic ibuprofen (IBU) on *C. jejuni* growth. Briefly, CPX was diluted in water to obtain an equally spaced gradient of 8 final concentrations, between 0 and 0.7 µM. For IBU, a 1:2 linear dilution was performed, to obtain a range of 8 final concentrations between 0 and 500 µM. The solutions were assembled in two-dimensions to obtain an 8×8 matrix of concentrations. The assay was performed in 96-well plates (Greiner BioOne), with *C. jejuni* precultures diluted to reach a starting OD_600_ = 0.001 in BB. Plates were sealed with breathable membranes (Breathe-Easy® sealing membrane, Merck) and incubated at 37°C in a microaerobic atmosphere, with shaking at ∼750 rpm. Growth was assessed by measuring OD_600_ with a plate reader (Infinite® M Plex, Tecan) and end-point growth across drug combinations was calculated for each well by subtracting OD_600_ values at 23 hours and at 1 hour.

### Time-kill assay

*C. jejuni* main cultures were grown in BB media until mid-exponential phase (OD_600_ ∼0.2-0.25), then split into subcultures of 5 ml in T25 flasks containing an equal volume of DMSO (untreated control) or 5xMIC of compounds, and incubated at 37°C in a microaerophilic atmosphere, with shaking. Samples were collected at hourly intervals, 10-fold serially diluted in BB in 96 well plates (Greiner BioOne) and 10 µL of each dilution spotted on 1x BB agar plates. Plates were incubated at 37°C in a microaerobic atmosphere for ∼40 hours or until detectable colonies appeared.

### Phenotypic analysis of compound treated *C. jejuni* by confocal microscopy

*C. jejuni* WT strain was grown and treated in the same conditions as for time-kill assay. At specific timepoints (1, 3, or 5 hours after treatment), 1 ml of bacteria was collected and centrifuged at 5,500 g for 5 min at RT. The pellets were resuspended in 500 µL 1X DPBS (Gibco™) and stained for DNA by adding 0.1 µL of Hoechst 33342 (16.2 mM, Thermo Fisher Scientific), and incubating with the dye for 10 min in the dark. An aliquot of 10 µL of the suspension was then mixed with 0.25 µL of the dye FM4-64 (5 mM, Thermo Fisher Scientific) to stain for membranes. After mixing, 1 µL of the resulting suspension was spotted on 8-well Ibidi µ-slides and covered with an agar pad (1% agarose in 1X DPBS) to immobilize bacteria before imaging. Microscopy images were immediately acquired using a Leica SP5 confocal laser scanning microscope.

### Membrane permeability assay (NPN assay)

A membrane permeability assay was performed as previously described ^40^, with minor modifications, to assess compound induced outer membrane damage. Briefly, WT *C. jejuni* was grown in BB medium until late exponential phase (OD_600_ ∼0.65), harvested by centrifugation at 5,500 x *g* for 5 min at RT, and resuspended in HEPES buffer (5 mM HEPES pH 7.4 + 5 mM glucose) to reach 2 OD/ml. For fluorescence measurements, 96 well black plates (Greiner) were added of 50 µL of the test compounds (at 4X the test concentration), 100 µL bacteria, and 50 µL of 40 µM 1-*N*-phenylnaphthylamine (NPN, Sigma) prepared in Hepes buffer. The plate was incubated at 37 °C for a total of 10 min with shaking. Fluorescence was assessed with a plate reader (Infinite® M Plex, Tecan; exc. 350 nm/em. 420nm). Background fluorescence of compounds was assessed by incubating 50 µL compounds with 150 µL HEPES buffer, while unspecific interaction of compounds with NPN was controlled for by incubating 50 µL of compounds, 100 µL Hepes buffer and 50 µL of 40 µM NPN, in the same conditions.

### RNA-seq

*C. jejuni* WT was grown under microaerobic conditions in 50 ml cultures in BB medium until mid-log phase (OD_600_ ∼ 0.25) and then split in 15 ml sub-cultures for 30 min treatment with 150 µM Ibuprofen (CAS 15687-27-1, Sigma) or DMSO. Cells were then harvested, mixed with 0.2 volumes of RNA Stop mix (5% buffer saturated phenol, pH 4.5 in ethanol), and immediately frozen in liquid N_2_. Samples were thawed on ice and cells pelleted at 4,500 x *g* for 20 min at 4°C. Total RNA was extracted from cell pellets using hot phenol as described previously (Sharma *et al.*, 2007). Total RNA was digested with *DNase*I and depleted of rRNA (RiboCop Meta, Lexogen). Libraries were prepared using an Adaptor Ligation protocol and sequenced on a NextSeq2000 instrument (100 cycles, single-end mode) at the Core Unit SysMed at the University of Würzburg. For RNA-seq data analysis, raw sequencing reads were subjected to quality and adapter trimming using Cutadapt ^71^ v5.0 with a cutoff Phred score of 20, and reads without any remaining bases were discarded (parameters: --nextseq-trim=20 -m 1 -a AGATCGGAAGAGCACACGTCTGAACTCCAGTCAC). Afterwards, all reads longer than 11 nt were aligned to the *C. jejuni* NCTC11168 reference genome (RefSeq assembly accession: GCF_000009085.1) using the READemption pipeline ^72^ v2.0.3 with segemehl ^73^ v0.3.4 and an accuracy cut-off of 95 % (parameters: -l 12 -a 95). Gene-wise read counts were quantified using READemption gene_quanti (parameters: --min_overlap 10 --features “5’-UTR,CDS,rRNA,tRNA,transcript,sRNA”), including previously determined 5’-UTRs and sRNA of *Campylobacter*. Based on these read counts, differential expression analysis was performed via DESeq2 ^74^ v1.42.1, using DESeq2 normalization (resulting in the following size factors: DMSO_r1/r2: 1.2414182, 0.8477668 and Ibu150_r1/r2: 1.1328194, 1.1012547). Shrinkage of log2FoldChange was performed using the apeglm package ^75^ v1.24.0 with default parameters. Significant features were selected based on thresholds for adjusted *p*-value after Benjamini-Hochberg correction *p*Adj < 0.05 and shrunken Log2FoldChange > 1.

### Text editing

Gemini was used to revise the initial draft of the manuscript text and for grammar checking.

## Acknowledgements

We thank Philipp Kible for help with cloning of *C. jejuni lux*-based transcriptional reporter strains and Sahil Sharma for cloning of CmeC epitope-tagged and CmeR deletion mutant strains. We thank all members of the Sharma lab for valuable scientific discussion. We thank the Core Unit Systems Medicine (University of Würzburg, GER) for support with RNA sequencing. This work was supported by a “StressRegNet” grant within the Bavarian Research Network (bayresq.net; www.bayresq.net) awarded to CMS, ARB and CLM.

## Authors contributions

The study was conceived by CMS, ARB, CLM, conceptualised by AZ, CMS, ARB, CLM, and experiments designed by AZ, CMS, CLM and ARB. AZ, MKA, and ARB established and optimized the high-throughput experiments. AZ performed high-throughput screening, follow-up and validation experiments. RW performed ligand docking, supervised by CS. TB supported the setup of fluorescence microscopy experiments. JK processed RNA-seq raw data. SLS established the use of luminescence reporters in *C. jejuni* and cloned the first *C. jejuni luxCDABE*-based transcriptional reporter strain. ROA, MF and AZ pre-processed screening data. AZ, ROA, an MF performed statistical analyses and data curation. All authors discussed and interpreted the data. AZ wrote the manuscript, with editing from MA, CMS, and ARB, and comments from all authors. CMS, ARB, CLM and CS secured funding and supervised this study. All authors approved the final version for publication.

## Data and code availability

To facilitate navigation of our dataset by the scientific community, we provide a user-friendly interactive application at (link). The data supporting the findings of this study will be available as source data & supplementary files. All code is available at (link). RNA-seq raw data, quantifications and results of the DESeq2 analysis will be deposited in the NCBI Gene Expression Omnibus (GEO) database.

