## Supplementary Figures for "Human-targeted drugs impact antibiotic sensitivity and growth of *Campylobacter jejuni*"

**This document contains:**

**Figures S1-S7**

**Supplementary References**

### Supplementary Figures

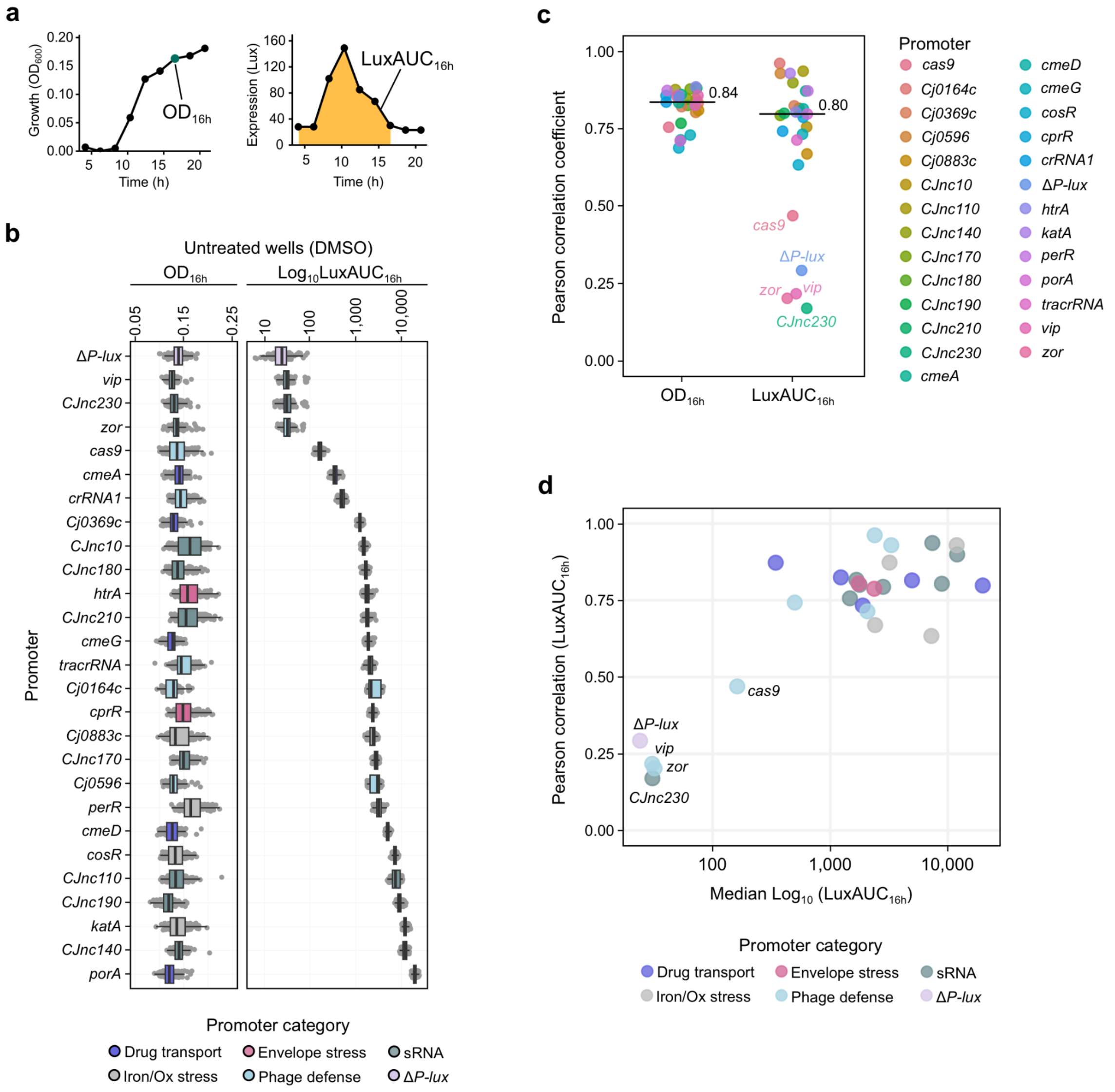


**Suppl. Fig. S1. Quality control and data analysis of the chemical genomics screen. a)** Example of baseline corrected growth and luminescence curves for the P*cmeA-lux* strain, in the presence of DMSO. Readouts for growth ($OD_{16h}$) and luminescence ($LuxAUC_{16h}$) and indicated. **b)** Boxplots of growth ($OD_{16h}$, left) and luminescence ($LuxAUC_{16h}$, right) across all reporters for untreated controls (DMSO wells). Each point represents the mean of two biological replicates for each DMSO well. Boxplots represent the distribution of mean replicate $OD_{16h}$/$LuxAUC_{16h}$values for each reporter strain. Center, upper, and bottom lines represent 25th, 50th, and 75th percentiles, whiskers extend to 1.5x inter-quartile range and points beyond whiskers are represented individually. $OD_{16h}$ = baseline-corrected OD600 at 16 hours; $LuxAUC_{16h}$ = area under the luminescence curve integrated till 16 hours. **c)** Pearson correlation of growth ($OD_{16h}$, left) and promoter activity ($LuxAUC_{16h}$, right) across two biological replicates depicted per reporter strain.

Median Pearson correlation coefficient across all reporter strains is indicated with a line and the exact value is given. **d)** Correlation between expression replicability and median promoter activity for all reporter strains. Scatterplot depicting the promoter-specific Pearson replicate correlation for promoter activity and the median promoter expression ($LuxAUC_{16h}$) for each reporter strain in DMSO wells.

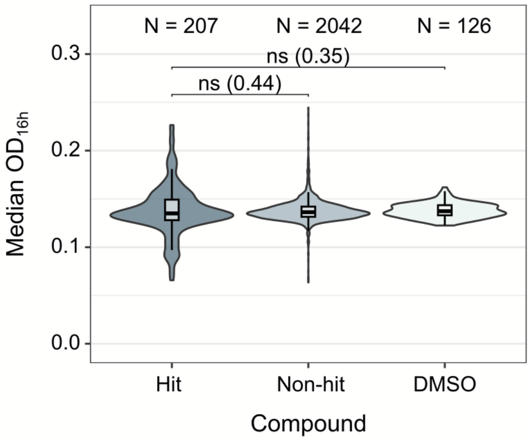


**Suppl. Fig. S2. Growth effects of hit and not-hit compounds.** Growth effects of the 2,249 compounds considered for CPI determination (excluding 166 compounds of the initial library of 2,415 small molecules, which displayed strong growth inhibitory effects). Hit and non-hit compounds were compared to DMSO controls, based on the median growth in the presence of the compound (optical density at 600 nm at 16 hours, $OD_{16h}$) across all strains. Results of Wilcoxon tests comparing the three different populations are indicated. Significance: ns = pAdj. > 0.05.

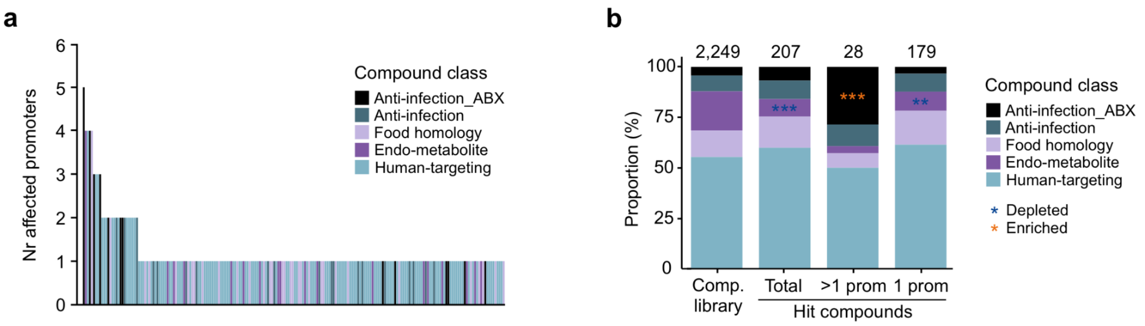


**Suppl. Fig. S3**. **Analysis of hit compounds. a)** Histogram showing the number of affected promoters for each hit compound, colored by class. **b)** Barplot showing the proportion of compound classes among the tested library (Comp. library), and the hits, including the total amount of compounds triggering at least one promoter (Total), more than one promoter (>1 prom) or one single promoter (1 prom). Results of Fisher's exact test for enrichment of compound classes among hit compounds compared to the screened library are indicated with asterisks. Orange = enriched, blue = depleted. Significance: *** = pAdj < 0.001, ** = pAdj <0.01.

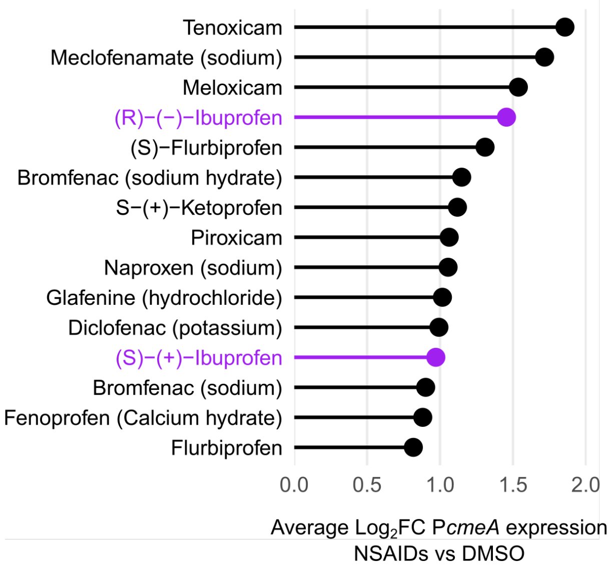


**Suppl. Fig. S4. NSAIDs activating P*cmeA* expression.** Average log2 fold-change (log2FC) of normalized expression of P*cmeA* in the presence of the respective compound, in comparison to DMSO.

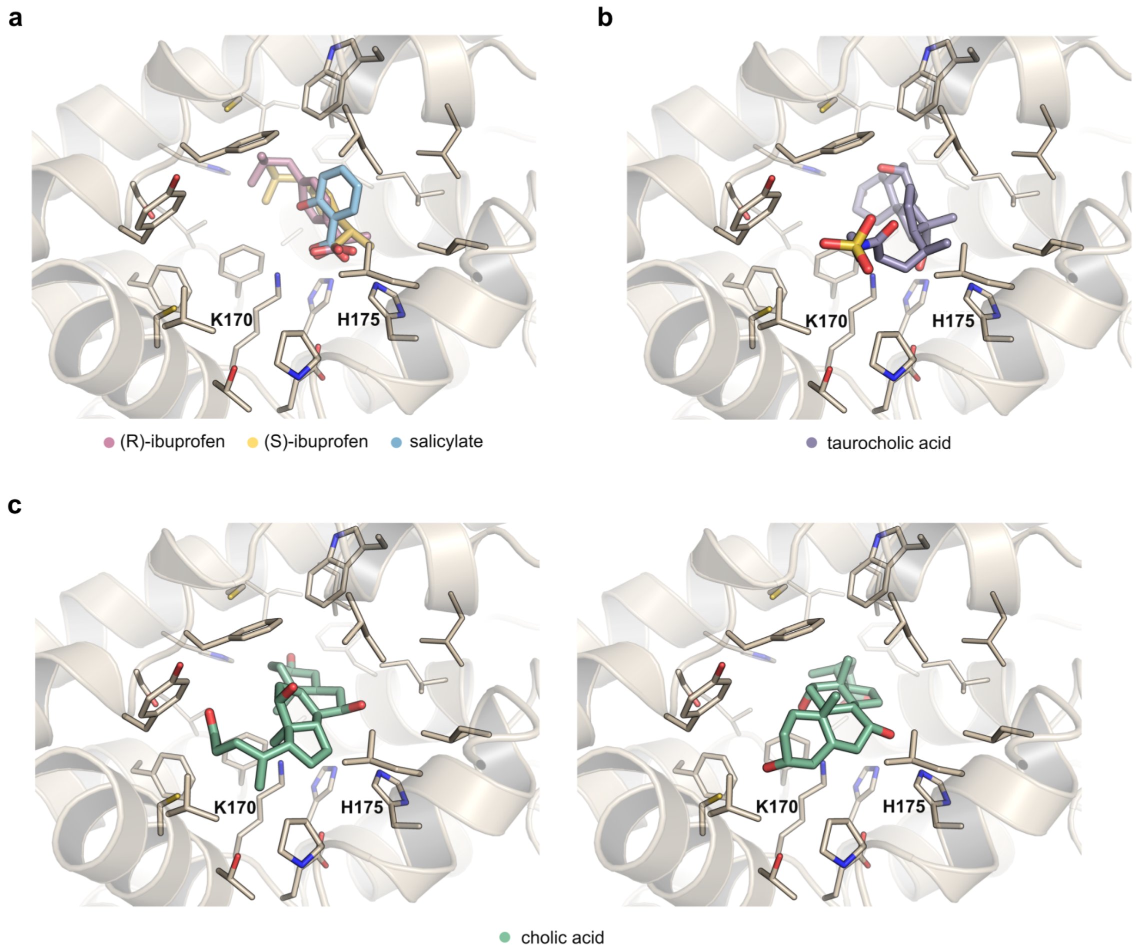


**Suppl. Fig S5. Binding modes of NSAIDs and bile salts in CmeR.** Comparison of the binding modes of the docked ligands (R/S)-ibuprofen and salicylate (a) and the bile salts in the available crystal structures: binding mode of taurocholic acid in PDB 3QQA (b) and binding modes of cholic acid as seen in PDB 3QPS (c, left) and PDB 3HGG (c, right) [1].

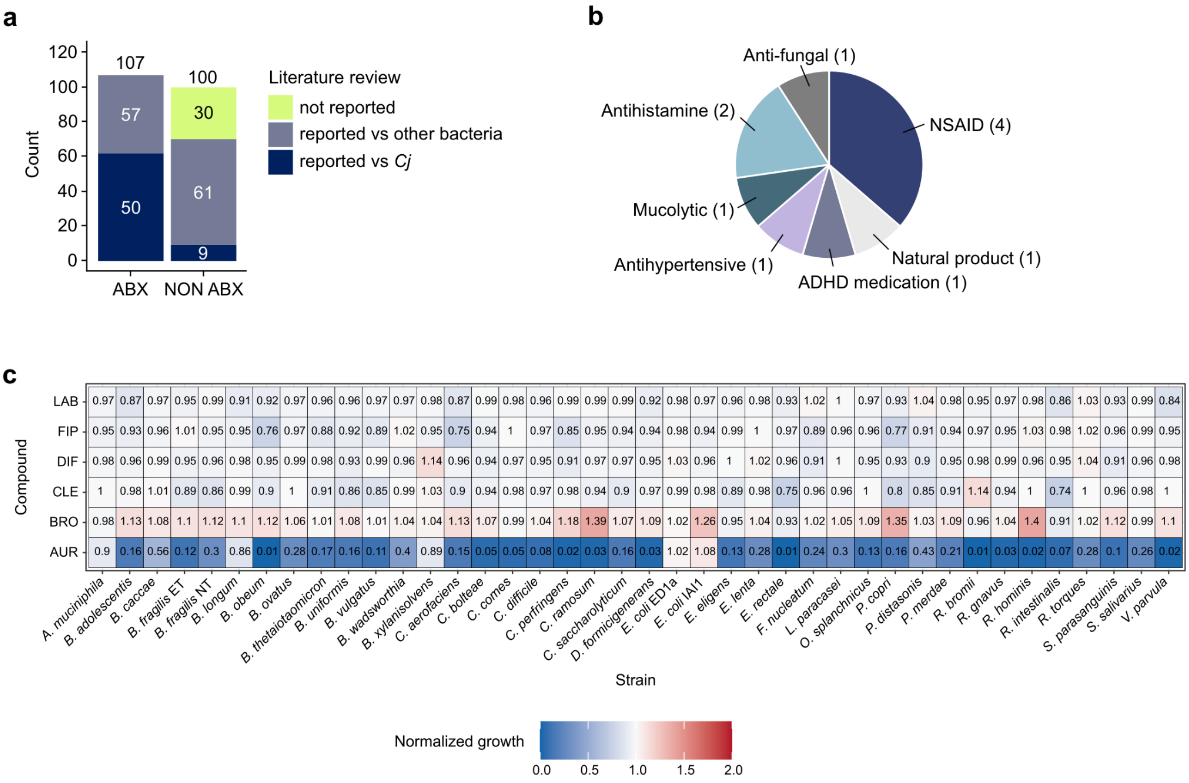


**Suppl. Fig. S6. Compounds inhibiting *C. jejuni* growth. a)** Literature review on growth inhibitors divided between antibiotics (ABX) and non-antibiotics (NON ABX), displaying reported growth inhibition against *Campylobacter* (Cj) or other bacteria (**Suppl. Table S8**). **b)** Proportion of therapeutic indications among *C. jejuni* growth inhibitors not displaying activity against 40 gut commensal species, according to [2] (n = 11; **Suppl. Table S8**). NSAID = Non Steroidal Anti Inflammatory Drugs, ADHD = Attention Deficit Hyperactivity Disorder. **c)** Selected *C. jejuni* growth inhibitors spare gut commensals. Heatmap showing the effect of selected human-targeted drugs (LAB = labetalol, FIP = fipexide, DIF = diflunisal, CLE = clemastine, BRO = bromhexine, AUR = auranofin) on the growth of human gut commensal species in the screen from Maier *et al.* [2]. Compound-treated growth, expressed as area under the growth curve (AUC), was normalized against inactive reference compounds. Auranofin is included as a representative human-targeted drug with broad-spectrum antibacterial activity against gut microbes (significant inhibition, adjusted $p$ < 0.01). Compounds from the *C. jejuni* screen selected for experimental validation showed no significant growth inhibition across human gut commensals (adjusted $p$ > 0.01).

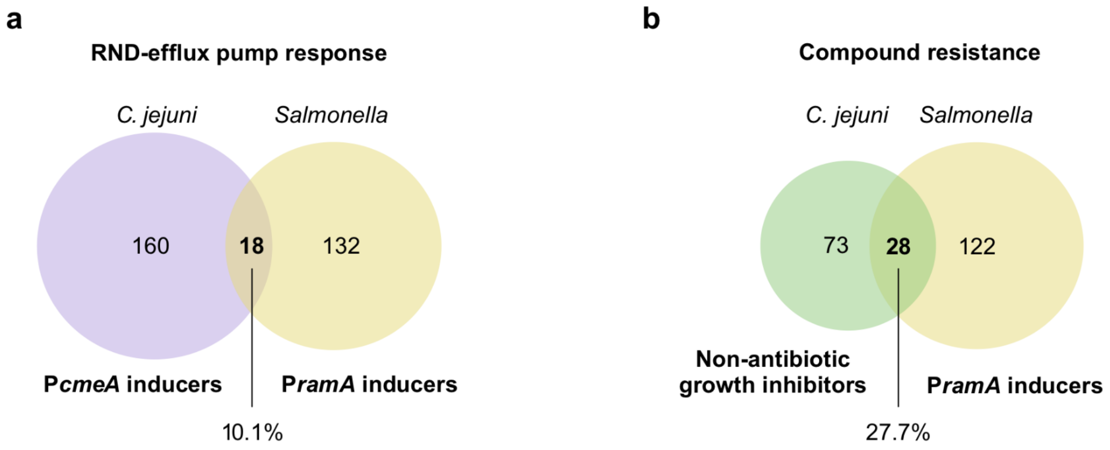


**Suppl. Fig. S7. Comparison between results of transcriptional chemical genomics screens performed in *C. jejuni* (this study) and *Salmonella* Typhimurium [3]. a)** Intersection of compounds activating the promoter of *C. jejuni* RND-efflux pump CmeABC (P*cmeA*) and of the AcrAB efflux pump activator RamA (P*ramA*) of *Salmonella*. **b)** Intersection between non-antibiotic compounds that inhibited *C. jejuni* growth and P*ramA* inducers. Percentages of compounds from the *C. jejuni* screen also identified in the *Salmonella* screen are displayed.
